# Early life glucocorticoid exposure modulates immune function in zebrafish (*Danio rerio*) larvae

**DOI:** 10.1101/2019.12.11.872903

**Authors:** Ruud van den Bos, Suzanne Cromwijk, Katharina Tschigg, Joep Althuizen, Jan Zethof, Robert Whelan, Gert Flik, Marcel Schaaf

**Affiliations:** Department of Animal Ecology and Physiology, Institute of Water and Wetland Research, Faculty of Science, Radboud University, Heyendaalseweg 135, 6525 AJ Nijmegen, the Netherlands; Animal Sciences and Health Cluster, Institute of Biology, Leiden University, 2333 CC Leiden, the Netherlands

**Keywords:** cortisol, glucocorticoid receptor, zebrafish, larvae, tail fin regeneration, lipopolysaccharide

## Abstract

In this study we have assessed the effects of increased cortisol levels during early embryonic development on immune function in zebrafish (*Danio rerio*) larvae. Fertilized eggs were exposed to either a cortisol-containing, a dexamethasone-containing (to stimulate the glucocorticoid receptor selectively) or a control medium for 6 hours post-fertilisation (0-6 hpf). First, we measured baseline expression of a number of immune-related genes (*socs3a*, *mpeg1.1*, *mpeg1.2* and *irg1l*) 5 days post-fertilisation (dpf) in larvae of the AB and TL strain to assess the effectiveness of our exposure procedure and potential strain differences. Cortisol and dexamethasone strongly up-regulated baseline expression of these genes independent of strain. The next series of experiments were therefore carried out in larvae of the AB strain only. We measured neutrophil/macrophage recruitment following tail fin amputation (performed at 3 dpf) and phenotypical changes as well as survival following LPS-induced sepsis (150 μg/ml; 4-5 dpf). Dexamethasone, but not cortisol, exposure at 0-6 hpf enhanced neutrophil recruitment 4 hours post tail fin amputation. Cortisol and dexamethasone exposure at 0-6 hpf led to a milder phenotype (e.g. less tail fin damage) and enhanced survival following LPS challenge compared to control exposure. Gene-expression analysis showed accompanying differences in transcript abundance of *tlr4bb, cxcr4a, myd88*, *il1β and il10*. These data show that early-life exposure to cortisol, which may be considered to be a model or proxy of maternal stress, induces an adaptive response to immune challenges, which seems mediated via the glucocorticoid receptor.

## 1. Introduction

In teleosts, like zebrafsh (*Danio rerio*), cortisol is the main endogenous corticosteroid, which is secreted when individuals perceive situations as stressful (Wendelaar Bonga, 1997). Like in other vertebrate species, cortisol binds in teleosts to the mineralocorticoid receptor (MR) and the glucocorticoid receptor (GR), which affect the transcription rates of genes following ligand binding (Alderman and Bernier, 2009; Alsop and Vijayan, 2008; Nesan and Vijayan, 2013). Since the GR has a lower affinity for cortisol than the MR, the GR mediates the actions of cortisol during stress, which involves optimizing energy expenditure by tuning the balance between and within physiological systems, like an organism’s metabolism and its immune, cardiovascular and central nervous system (e.g. Gorissen and Flik, 2016; Nesan and Vijayan, 2013; Wendelaar Bonga, 1997). Following long-term exposure to stress, baseline levels of cortisol are increased, reflecting the allostatic load that the environment imposes on an organism (Gorissen and Flik, 2016; McEwen and Wingfield, 2003). Cortisol has been shown to signal through the GR already during the very early stages of embryonic development; in oocytes maternally deposited cortisol and GR mRNA are present (Pikulkaew *et al*., 2010, 2011; Wilson *et al*., 2013). These cortisol levels may reflect the allostatic load that the mothers experience in their environment and it has therefore been hypothesized that these deposited cortisol levels are important for preparing the offspring for the expected allostatic load that larvae will encounter in the prevailing environment, thereby programming their cortisol secretion and the functioning of physiological systems to meet expected demands (Nesan and Vijayan, 2013; van den Bos *et al*., 2019). While in previous studies we focussed on the effects of cortisol exposure (between 0 and 6 hours post fertilisation (hpf)) on vigilance-related behaviour, baseline cortisol levels and metabolism in larvae (van den Bos *et al*., 2019; van den Bos, 2019) in the present study we focussed on the functioning of the immune system, in particular the innate immune system on which zebrafish larvae rely (Meijer and Spaink, 2011). Cortisol exposure may be considered to be a model or proxy of maternal stress (Best *et al*., 2017; Nesan and Vijayan, 2012, 2016; van den Bos *et al*., 2019).

Zebrafish is a highly suitable animal model to study early life events in the fields of biomedical research, behavioural biology and eco-toxicology (e.g. Champagne *et al*., 2010; Dai *et al*., 2013; Nesan and Vijayan, 2013; Steenbergen *et al*., 2011; Stewart *et al*., 2014). Fertilized eggs develop into independently feeding larvae outside the mother, without parental care, and can easily be maintained under different experimental conditions as well as pharmacologically manipulated. In zebrafish, it has been demonstrated that mothers deposit cortisol and GR mRNA in oocytes (Pikulkaew *et al*., 2010, 2011; Wilson *et al*., 2013). These cortisol levels decrease over the first 24 hours post fertilisation, after which zygotes start to produce cortisol by the then developing interrenal cells (Liu, 2007; Pikulkaew *et al*., 2010, 2011; Wilson *et al*., 2013). After hatching (48-72 hpf) pituitary control over interrenal cortisol production starts and it takes another 4-5 days before the hypothalamus-pituitary-interrenal (HPI) axis is fully functionally mature (Alderman and Bernier, 2009; Alsop and Vijayan, 2008). Maternal GR mRNA is present during the first 6 hpf, and at 8-9 hpf zygotic expression of the GR commences, while the MR mRNA production starts at 24 hpf (Alsop and Vijayan, 2008; Pikulkaew *et al*., 2010, 2011).

In several studies the effect of cortisol exposure during early embryonic stages has been investigated in zebrafish. In these studies cortisol levels were increased by injection of cortisol into the yolk of one-cell stage embryos (e.g. Best *et al*., 2017; Nesan and Vijayan, 2012, 2016) or through addition of cortisol to the medium (Hartig *et al*., 2016; van den Bos *et al*., 2019). These studies showed that, as a result of the cortisol exposure during early embryonic stages, larval baseline levels of cortisol were increased (Best *et al*., 2017; Hartig *et al*., 2016; Nesan and Vijayan, 2012; van den Bos *et al*., 2019). For example, in a recent study, we have demonstrated that cortisol exposure between 0 and 6 hpf increased baseline cortisol levels 5 days post fertilisation (dpf) and this effect was stronger in larvae from the AB strain than in larvae from the TL strain (van den Bos *et al*., 2019).

Exposure to cortisol (0-5 dpf) has been shown to lead to an enhanced expression of immune-related genes in zebrafish larvae at 5 dpf (Hartig *et al*, 2016), suggesting that early cortisol exposure increases the activity of the immune system. In the present study, we have first measured the expression of a selected number of these up-regulated genes at 5 dpf (*socs3a*, *mpeg1.1*, *mpeg1.2*, *irg1l*) following 0-6 hpf exposure to cortisol in zebrafish larvae of the AB strain to assess whether our method produces similar effects. Tüpfel long-fin (TL) is another widely used zebrafish strain next to AB and is characterized by spots rather than stripes as well as long fins rather than short fins. In previous studies we have observed consistent differences between larvae of the AB strain and larvae of the TL strain at the level of both HPI-axis activity and behaviour (van den Bos *et al*., 2017a, 2017b, 2019a, 2020). We have attributed these differences to the mutation in the connexin 41.8 gene that leads to spots (for discussion: see van den Bos *et al*., 2020). Measuring the expression of these genes in larvae of the TL strain next to larvae in the AB strain may therefore reveal how robust our findings are. Finally, to assess the role of the GR in more detail, we exposed fertilized eggs 0-6 hpf to dexamethasone, a specific GR agonist (Rupprecht *et al*., 1993).

To functionally assess the activity of the immune system following early life exposure to cortisol or dexamethasone, we used two experimental models for immune activation. First, we used the tail fin amputation assay. This is a well-established model in which amputation of the tail triggers expression of many pro-inflammatory molecules and the recruitment of innate immune cells (neutrophils and macrophages) towards the wounded area (Hall *et al*., 2014; Renshaw *et al*., 2006; Roehl, 2018). The tail fin amputation assay was performed using the double transgenic fish line *Tg(mpx:GFP/mpeg1:mCherry-F)* (Bernut *et al*., 2014; Renshaw *et al*., 2006). Recruitment of neutrophils and macrophages was determined following tail fin amputation in larvae at 3 dpf (Chatzopoulou *et al.,* 2016; Xie *et al*., 2019). Second, we used a sepsis model, which involved a challenge with lipopolysaccharide (LPS), the membrane component of Gram-negative bacteria, in 4 dpf larvae. We measured survival, phenotypical changes, and the expression of a series of LPS-responsive genes (Dios *et al*., 2014; Hsu *et al*., 2018; Novoa *et al*., 2009; Philip *et al*., 2017).

## 2. Materials and Methods

### 2.1. Subjects, spawning and care

Housing conditions and breeding procedures were similar as those reported in van den Bos *et al*. (2017a, 2017b, 2019; AB and Tüpfel long-fin (TL) strains) or Xie *et al*. (2019; the double transgenic fish line *Tg(mpx:GFP/mpeg1:mCherry-F)*. They were kept in recirculation systems (~28°C) under a 14h:10h light-dark cycle and fed twice daily.

Breeding started at least one hour after the last feeding of zebrafish (>16:00 h). Males and females of the AB or TL strain were placed in a zebrafish breeding tank, separated by a partitioning wall, with water of ~28°C. After turning on the lights the next morning, the partitioning wall was removed and tanks were placed at a slight angle, such that the fish had the possibility to move into shallow water to spawn.

### 2.2. Cortisol/dexamethasone exposure during early embryonic development (0-6 hpf)

Cortisol (hydrocortisone; Sigma-Aldrich, Zwijndrecht, the Netherlands) and dexamethasone (Sigma-Aldrich, Zwijndrecht, the Netherlands) were dissolved in 96% ethanol in the required stock solution concentrations and stored at −20°C. From these stock solutions media with the appropriate concentration were freshly prepared for each experiment (Althuizen, 2018; van den Bos *et al*., 2019): cortisol-containing medium: 400 µg/l cortisol (1.1 µM), 0.4 ml/l 96% ethanol (0.04% v/v), 5 mM NaCl, 0.17 mM KCl, 0.33 mM CaCl_2_, 0.33 mM MgSO_4_ in dH2O; dexamethasone-containing medium: 430 µg/l dexamethasone (1.1 µM), 0.4 ml/l 96% ethanol (or 1 ml/l 96% ethanol (0.1% v/v)); depending on the specific experiment), 5 mM NaCl, 0.17 mM KCl, 0.33 mM CaCl_2_, 0.33 mM MgSO_4_ in dH2O. Control medium consisted of: 0.4 ml/l 96% ethanol (or 1 ml/l 96% ethanol; depending on the specific experiment), 5 mM NaCl, 0.17 mM KCl, 0.33 mM CaCl_2_, 0.33 mM MgSO_4_ in dH2O.

Directly following spawning and fertilization, eggs were collected and randomly assigned to Petri dishes filled with either cortisol-containing, dexamethasone-containing or control medium. Within 1-1.5 hpf Petri dishes were placed in an incubator set at 28.5 °C (300–350 lux). Eggs were exposed to these solutions for 6 hrs. It has been shown that both cortisol and dexamethasone diffuse inside the eggs in this period (Steenbergen *et al*., 2017). In addition, we have shown this procedure to be effective in eliciting changes in physiology and behaviour at 5 dpf (Althuizen, 2018; van den Bos *et al*., 2019).

Following this, cortisol-containing, dexamethasone-containing and control media were replaced by E3 medium (5 mM NaCl, 0.17 mM KCl, 0.33 mM CaCl_2_, 0.33 mM MgSO_4_, 3 ml/l 0.01% (w/v) methylene blue in dH2O). Embryos were rinsed twice in E3 medium to ensure that the original treatment media were completely removed. Petri dishes were returned to the incubator allowing the embryos to develop further (28.5°C; 14h:10h light-dark period (lights on: 09.00h – 23.00h); light phase: 300–350 lux; dark phase; 0 lux). At 1 and 4 dpf E3 medium was refreshed and unfertilized eggs, dead eggs/embryos/larvae and chorions were removed from the dishes.

All experiments were carried out in accordance with the Dutch Experiments on Animals Act (http://wetten.overheid.nl/BWBR0003081/2014-12-18), the European guidelines for animal experiments (Directive 2010/63/EU; http://eur-lex.europa.eu/legal-content/NL/TXT/HTML/?uri=CELEX:32010L0063) and institutional regulations (Radboud University or Leiden University). Larvae were euthanized by placing them in ice slurry for at least 20 minutes followed by adding bleach to the slurry. In case of anaesthesia 0.01% 2-phenoxyethanol (Sigma Aldrich, Zwijndrecht, the Netherlands; experiments Radboud University) or 0.02% buffered aminobenzoic acid ethyl ester (tricaine; Sigma Aldrich, Zwijndrecht, the Netherlands; experiments Leiden University) was used.

### 2.3 Baseline gene expression analysis

Following 0-6 hpf exposure to the different media, larvae of the AB and TL strain were sampled for gene expression at 5 dpf between 16:00 hrs and 19:00 hrs (van den Bos *et al*., 2019; experimental time-line: figure 1). Genes of interest were: *socs3a*, *mpeg1.1*, *mpeg1.2* and *irg1l* (for primer sequences of these genes: see van den Bos *et al*., 2017a). Gene expression was determined by qPCR analysis as described below.

**Figure 1:**
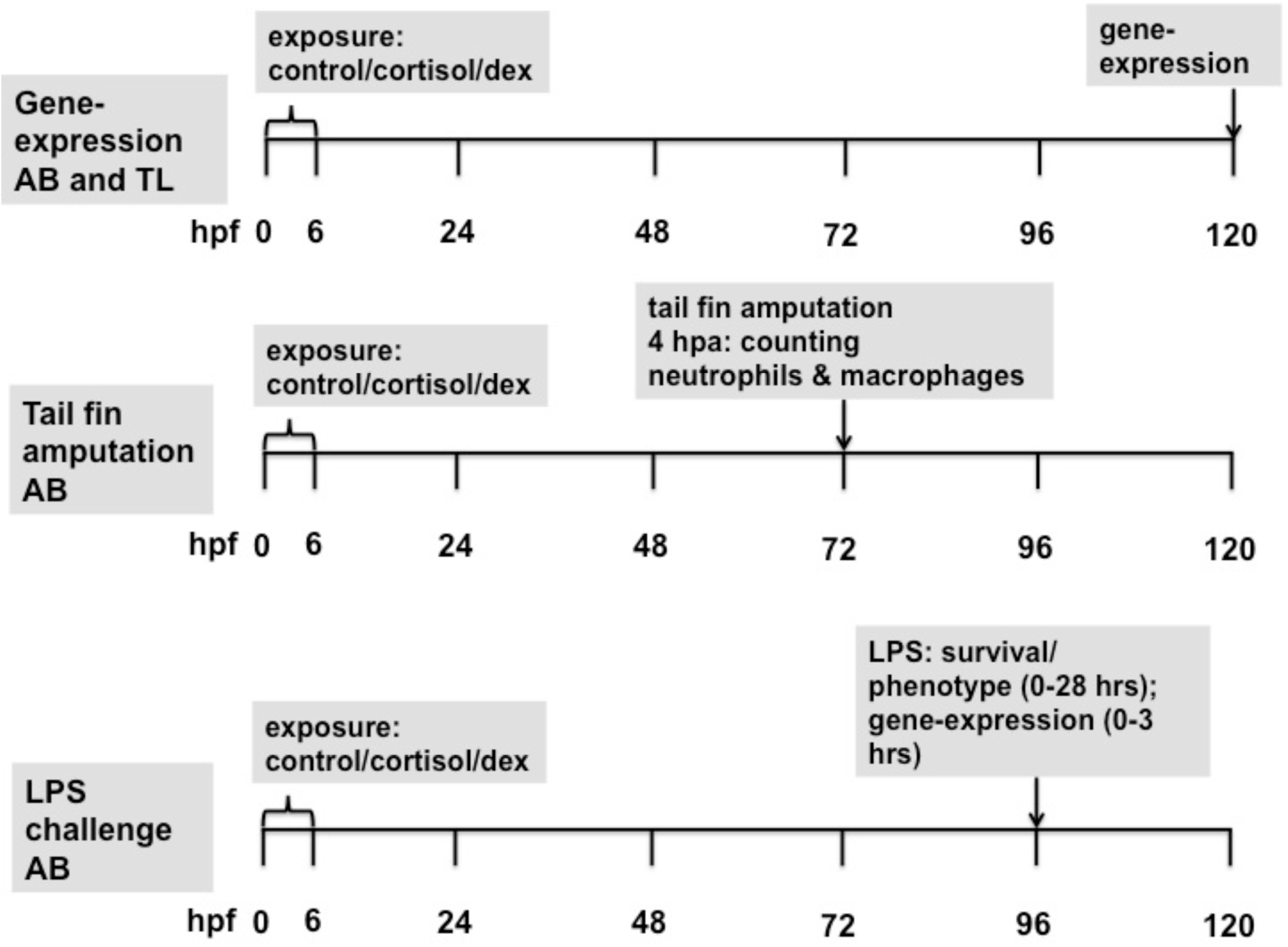
Experimental time-line of the different experiments. dex=dexamethasone

### 2.4 Tail fin amputation assay

The tail fin amputation assay was performed using the double transgenic fish line *Tg(mpx:GFP/mpeg1:mCherry-F;* Bernut *et al*., 2014; Renshaw *et al*., 2006). Three-day-old zebrafish larvae were anaesthetized in E3 medium containing 0.02% buffered tricaine and loaded onto 2% agarose-coated Petri dishes (experimental time-line: figure 1). Amputation was performed with a 1 mm sapphire blade (World Precision Instruments, Friedberg, Germany) using a Leica M165C stereomicroscope and a micromanipulator (Leica Microsystems BV, Amsterdam, the Netherlands; see below, figure 3A). Larvae were fixed in 4% paraformaldehyde in PBS at 4 hours post amputation (hpa) and stored overnight at 4°C. The following day larvae were washed twice for one minute and then twice for five minutes in PBS containing 0.01% Tween 20 (Sigma Aldrich, Zwijndrecht, the Netherlands).

A Leica M205FA fluorescence stereomicroscope supported by LAS software (version 4.12.0; Leica Microsystems BV, Amsterdam, the Netherlands) was utilised to visualise the leukocytes. Detection of neutrophils and macrophages was based on their fluorescent GFP and mCherry signals respectively. To quantify cell migration towards the wounded area, cells within a distance of 200µm from the amputation site were counted manually, as previously described (Xie *et al*., 2019). Data (numbers of migrated neutrophils and macrophages per individual) were pooled from three individual experiments (n>10 per experiment), and the presented data are means (± SEM).

### 2.5. LPS exposure

First, we conducted a pilot study to assess the optimal LPS dose, exposure duration and parameters to be measured (protocols adapted from: Dios *et al*., 2014; Hsu *et al*., 2018; Novoa *et al*., 2009; Philip *et al*., 2017). Incubation for 30 minutes in 150 μg/ml LPS (B11.04; Sigma Aldrich, Zwijndrecht, the Netherlands) was effective in eliciting a robust increase in *il1β* expression (assessed using qPCR analysis), changes in tail fin morphology (swollen or damaged tails) and increased levels of reactive oxygen species (ROS; measured by a fluorescent labelling method according to Philip *et al*., 2017). Hence, we used this dose in subsequent experiments.

Subsequently, two experiments were conducted. In both experiments 4 dpf larvae were exposed to 150 μg/ml LPS (B11.04; Sigma Aldrich, Zwijndrecht, the Netherlands) in E2 medium or control E2 medium for 30 minutes in Petri dishes (n=50 in 25 ml; experimental time-line: figure 1). Following this exposure, LPS-containing medium or control E2 medium was replaced by fresh E2 medium (larvae were rinsed two times to ensure that the original media were removed). Larvae either remained in the Petri dishes for sampling for gene expression at later time points or were transferred individually to 24 wells plates (Greiner Bio-One BV, Alphen a/d Rijn, the Netherlands) for assessing phenotypical changes and survival (volume per well: 1-1.5 ml). Six treatment groups were thus created: 0-6 hpf control, cortisol or dexamethasone treatment, combined with either 4 dpf LPS or control treatment (for 30 min).

In the first experiment the level of gene expression at 0 hr, 0.5 hr (i.e. directly following exposure), 1 hr (i.e. 30 minutes after ending exposure) and 3 hrs (i.e. 2.5 hrs after ending exposure; Novoa *et al*., 2009) was determined by qPCR analysis as described below. Genes of interest (see papers by: Hsu *et al*., 2018; Kanwal *et al*., 2013; Novoa *et al*., 2009; Philip *et al*., 2017) were genes encoding proteins involved in barrier function of the vascular endothelium (*cldn5a*, *cldn2*, *oclnb*), Toll-like receptors (*tlr2*, *tlr4ba*, *tlr4bb*, *tlr5a*, t*lr5b*), and regulators of the immune response (*il1β*, *il10*, *myd88*, *cxcr4a, cxcr4b, ptpn6*). Primer sequences are listed in Table 1.

**Table 1:**
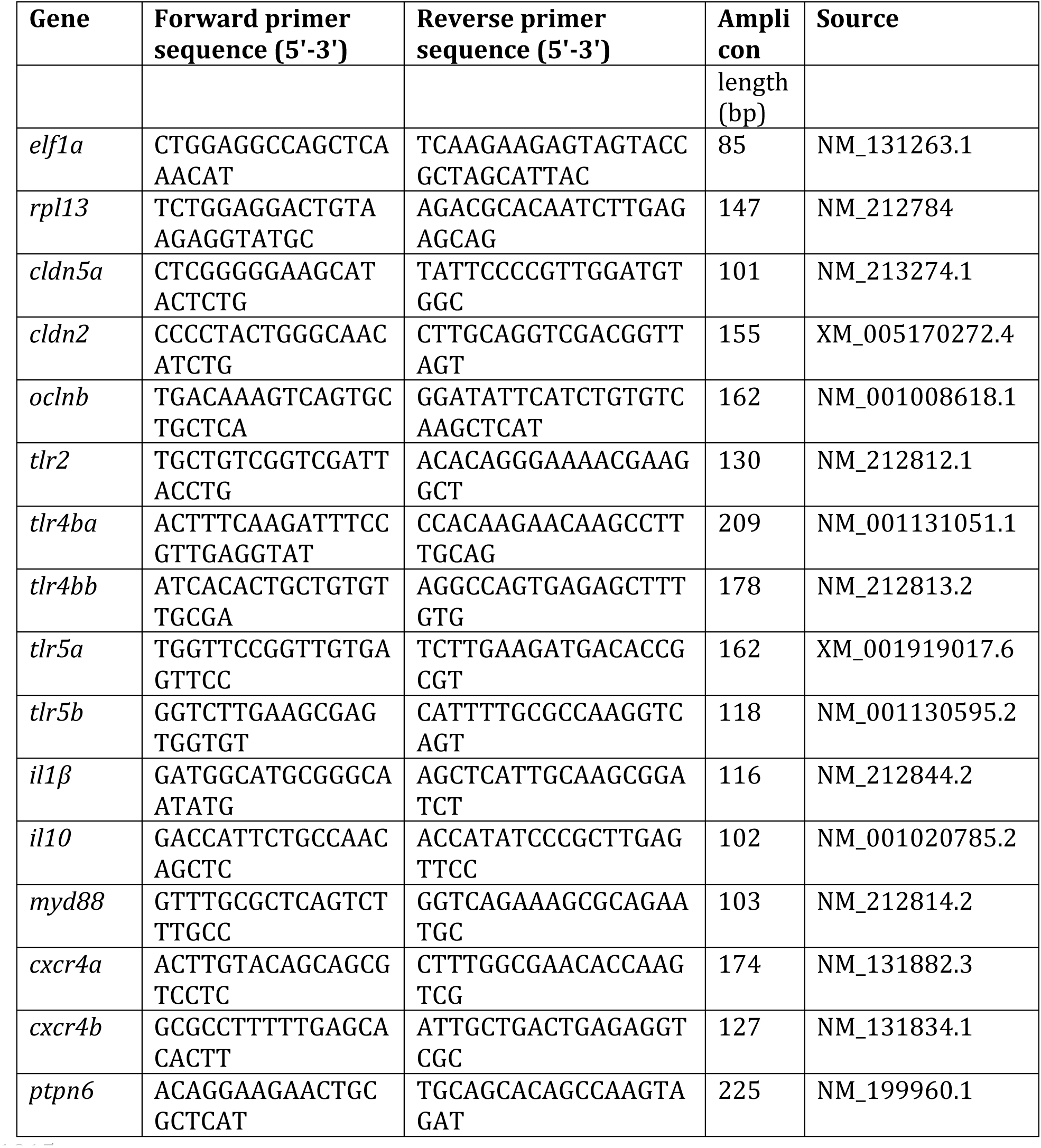
Nucleotide sequences of forward and reverse primers used for qPCR. Reference genes: elongation factor 1α (*elf1α*); ribosomal protein L13 (*rpl13*)

In addition, in the first experiment survival and phenotypical changes were determined at 0, 4 (i.e. 3.5 hrs after ending exposure) and 28 hrs (i.e. 27.5 hrs after ending exposure). Phenotypical changes included (see Philip *et al*., 2017): changes in tail fin morphology (normal, swollen (oedema) or damaged), presence of heart oedema and changes in shape (straight or curved). In the second experiment (that also served as replicate for the first experiment) survival and phenotypical changes were measured at 0, 0.5, 1, 3, 6 and 24 hrs (i.e. before exposure, directly after exposure, 0.5 hrs after ending exposure, 2.5 hrs after ending exposure, 5.5 hrs after ending exposure and 23.5 hrs after ending exposure).

### 2.6 Gene expression analysis

For the assessment of gene expression levels by qPCR analysis, 3-5 larvae were transferred to a 2-ml Eppendorf tube; thus, one sample contained material from 3-5 larvae. Residual medium was removed with a pipette, tubes were snap frozen in liquid nitrogen, kept on ice during the sampling procedure, and subsequently stored at −80 °C until total RNA extraction.

RNA isolation, RNA preparation, removal of genomic DNA from the samples and synthesis of cDNA was performed according to the protocol described in van den Bos *et al*. (2017a). Total RNA content of each sample was isolated. This was done by homogenising the tissue with 400 µl Trizol reagent (Invitrogen, Carlsbad, USA) in a Grinding Mill (Retsch GmbH, Germany) for 20 s at 20 Hz. After homogenisation, samples were kept at room temperature for 5 min. Next, 80 µl chloroform was added and the solution was mixed by shaking for 15 s. Afterwards, samples were kept at room temperature for 2 min. The samples were centrifuged at 14,000 rpm for 10 min in a cooled centrifuge (4°C) and the aqueous phase of the samples was transferred to a new tube. To this phase, 200 µl isopropanol was added and this solution was mixed well by inversion of the tube. The solution was then stored at −20°C for 2 h and centrifuged afterwards for 15 min at 14,000 rpm in a cooled centrifuge (4°C). The supernatant was decanted and the pellet washed with 500 µl 75% ethanol and centrifuged 10 min at 14,000 rpm in a cooled centrifuge (4°C). The supernatant was decanted, after which the pellet was centrifuged for 5 s to remove all the remaining supernatant using a pipette. The pellet containing the RNA was air-dried for 10 min at room temperature and afterwards dissolved in 100 µl ice cold DEPC-treated dH2O. To this RNA solution, 10 µl 3M NaAc (pH 5.4) and 250 µl 100% ethanol were added. The solution was mixed by inverting the tube and samples were stored for 2 h at −20°C. Subsequently, the samples were centrifuged for 15 min at 14,000 rpm in a cooled centrifuge (4°C), and the supernatant was decanted and the pellet washed washed as described earlier. Finally, the RNA pellet was dissolved in 15 µl DEPC-treated dH2O. The concentration and quality of RNA in each sample were measured using a nanodrop spectrophotometer at 260 nm wavelength (Nanodrop, Wilmington, DE, USA).

Isolated RNA was treated with DNase to remove any (genomic) DNA from the sample; 400 ng RNA was transferred into a PCR strip, and DEPC-treated dH2O was added to a volume of 8 µl. To this, 2 µl of DNase mix was added, containing 1 µl 10x DNase I reaction buffer and 1 µl (1 U/µl) amplification grade DNase I (both from Invitrogen, Carlsbad, USA). The resulting mix was incubated for 15 min at room temperature. Afterwards, 1 µl 25 mM EDTA was added to stop the DNase reaction and the reaction mix was incubated for 10 min at 65°C and returned on ice.

After the DNase treatment, samples were used to synthesize cDNA by the addition of 1 µl random primers (250 ng/µl), 1 µl 10 mM dNTP mix, 4 µl 5 x 1st strand buffer, 1 µl 0.1M DTT, 1 µl RNase inhibitor (10 U/µl), 0.5 µl Superscript II (reverse transcriptase) (200 U/µl) (all from Invitrogen, Carlsbad, USA) and 0.5 µl DEPC-treated dH2O. The resulting mix was incubated for 10 min at 25°C for annealing of the primers and then 50 min at 42°C for reverse transcription. Enzymes were hereafter inactivated by incubating samples at 70°C for 15 min. Finally, 80 µl dH2O was added to dilute the samples five times for the qPCR reaction.

To measure the relative gene expression in each sample, real-time qPCR was carried out for each gene of interest. For each qPCR reaction, 16 µl PCR mix (containing 10 µl SYBR green mix (2x) (BioRad, Hercules, USA), 0.6 µl forward and reverse gene-specific primer (10 µM) and 4.8 µl H2O) was added to 4 µl of cDNA. The qPCR reaction (3 min 95°C, 40 cycles of 15 s 95°C and 1 min 60°C) was carried out using a CFX 96 (BioRad, Hercules, USA) qPCR machine. Analysis of the data was carried out using a normalisation index of two reference genes (viz. *elongation factor alpha* (*elf1a*) and *ribosomal protein L13* (*rpl13*)) (Vandesompele *et al*., 2002).

### 2.7. Statistics

For gene expression analyses, outliers were removed following Grubb’s outlier test (p≤0.01). We explored the interrelationships of transcript abundance levels using Principal Component Analysis (PCA) with orthogonal rotation (Varimax rotation with Kaiser normalization; see van den Bos *et al*., 2017b). In case of missing samples, data were excluded list-wise. The number of retained components was based on eigenvalues (>1) and visual inspection of the scree plot. The Kaiser-Meyer-Olkin (KMO) measure of sampling adequacy and Bartlett’s test of sphericity were done to ensure that data obeyed analysis criteria; both are measures to assess whether the correlation matrix is suited for factor analysis (Budaev, 2010). Component scores were saved and used for further statistical analysis. The following component loading cut-off points were considered: ≤−0.600 or ≥0.600 (Ferguson, 1989; Budaev, 2010).

For the basal gene expression values a two-way or three-way Analysis of Variance (ANOVA) was run with treatment, strain or batch (where applicable) as independent factors. In the tail fin amputation assay a Student’s t-test was run on the number of neutrophils or macrophages at 4 hpa comparing 0-6 hpf treatment groups (cortisol *versus* control; dexamethasone *versus* control). In the LPS exposure experiment, for gene expression a multivariate analysis of variance (MANOVA) was run (to account for multiple comparisons) followed by univariate analysis of variance (0-6 hpf treatment and time as independent factors followed by post-hoc testing (Tukey HSD)). In addition per time point a one-way ANOVA was run with 0-6 hpf treatment as factor followed by post-hoc testing (Tukey HSD).

In the LPS exposure experiment differences in survival rate were assessed using the Kaplan-Meijer procedure (Log Rank Mantel-Cox). Differences in phenotypical changes were compared using Chi-square tests.

Significance was set at p≤0.05 and trends are indicated (p≤0.10) where appropriate; ns: not significant: p>0.10. Unless otherwise stated all p-values are two-tailed. All statistical analyses were done using IBM SPSS version 23 (IBM, Armonk, NY, USA).

## 3 Results

### 3.1 Baseline gene expression analysis

Rather than analysing transcript abundance of different genes following glucocorticoid treatment (cortisol or dexamethasone, 0-6 hpf) separately, we explored the effects of exposure through the interrelationships of transcript abundance of genes using PCA.

A PCA for the 0-6 hpf cortisol treatment experiment revealed two components (figure 2A; supplementary figure 1A shows the transcript abundance of individual genes). The KMO value was sufficiently high (0.522) and Bartlett’s test of sphericity was highly significant (Chi-square=17.707, df=6, p≤0.007), indicating that the data were adequate for a PCA. The first component (explaining 44.0% of the variance) was comprised of *irg1l (*loading: 0.854) and *socs3a* (loading: 0.890); the second component (explaining 28.9% of the variance) of *mpeg1.1* (loading: 0.890) and *mpeg1.2* (loading: 0.890). Each component was analysed separately using ANOVA. Treatment (0-6 hpf) with cortisol enhanced transcript abundance of genes in component 1 (*irg1l* and *socs3a*) independent of strain (three-way ANOVA (strain, treatment, batch): treatment: F(1,23)=6.477, p≤0.018). Transcript abundance was overall higher in TL than AB larvae (strain: F(1,23)=7.023, p≤0.014). Treatment (0-6 hpf) with cortisol enhanced transcript abundance of genes in component 2 (*mpeg1.1* and *mpeg1.2)* independent of strain (three-way ANOVA (strain, treatment, batch): treatment: F(1,23)=7.660, p≤0.011).

**Figure 2.**
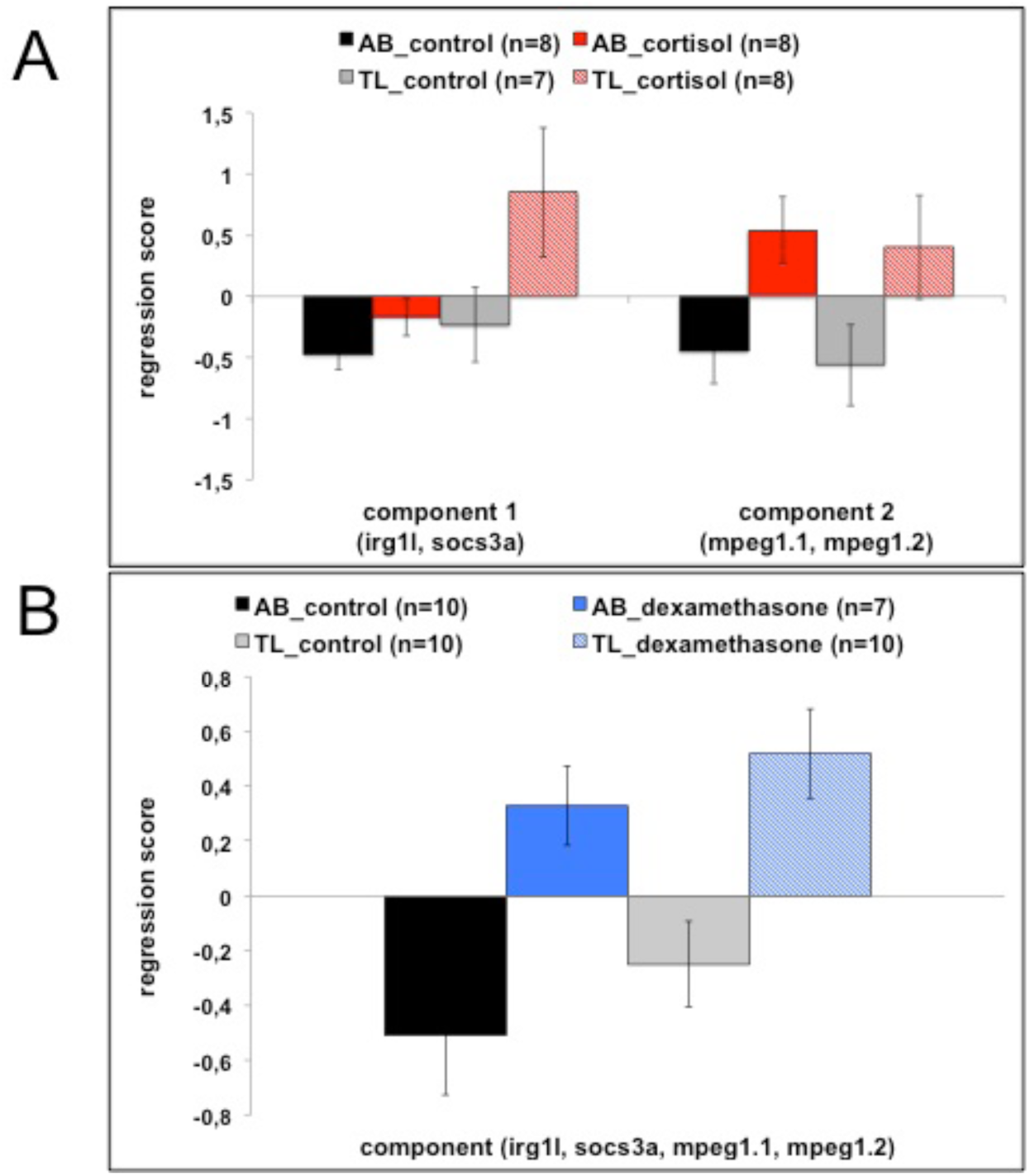
**panel A**: Regression scores (mean± SEM) of components 1 and 2 from the PCA for the different treatments (cortisol or control) and strains (AB or TL). Genes that contributed to the components are indicated in the figure. One subject was removed from the statistical analyses (TL control) as it was a consistent outlier following Grubb’s outlier test. **panel B**: Regression scores (mean± SEM) of the only component of the PCA for the different treatments (dexamethasone or control) and strains (AB or TL). Genes that contributed to the component are indicated in the figure.

A PCA for the 0-6 hpf dexamethasone treatment experiment revealed only one component (figure 2B; supplementary figure 1B shows the transcript abundance of individual genes). The KMO was sufficiently high (0.626) and Bartlett’s test of sphericity was highly significant (Chi-square=102.945, df=6, p<0.001), indicating that the data were adequate for a PCA. This component explained 76.6% of the variance. Loadings onto this component were: *irg1l (*0.867). *socs3a* (0.835), *mpeg1.1* (0.928) and *mpeg1.2* (0.868). Treatment with dexamethasone enhanced transcript abundance of genes independent of strain (two-way ANOVA (strain, treatment): treatment: F(1,33)=6.745, p≤0.014).

These data show that glucocorticoid treatment was effective in eliciting changes in baseline expression of (a selected set of) immune-related genes.

### 3.2 Tail fin amputation assay

Figure 3A shows the site of the tail fin amputation performed at 3 dpf. Neutrophil recruitment at 4 hpa was enhanced in the 0-6 hpf dexamethasone treatment group (Student’s t-test: t=2.917, df=106, p≤0.004; figure 3C), but not in the 0-6 hpf cortisol treatment group (Student’s t-test: t=0.621, df=89, ns; figure 3B), compared to the group treated with control medium at 0-6 hpf. No effects were found for macrophage recruitment for either 0-6 hpf cortisol treatment (Student’s t-test: t=0.923, df=89, ns; figure 3B) or 0-6 hpf dexamethasone treatment (Student’s t-test: t=-1.095, df=106, ns; figure 3C).

**Figure 3.**
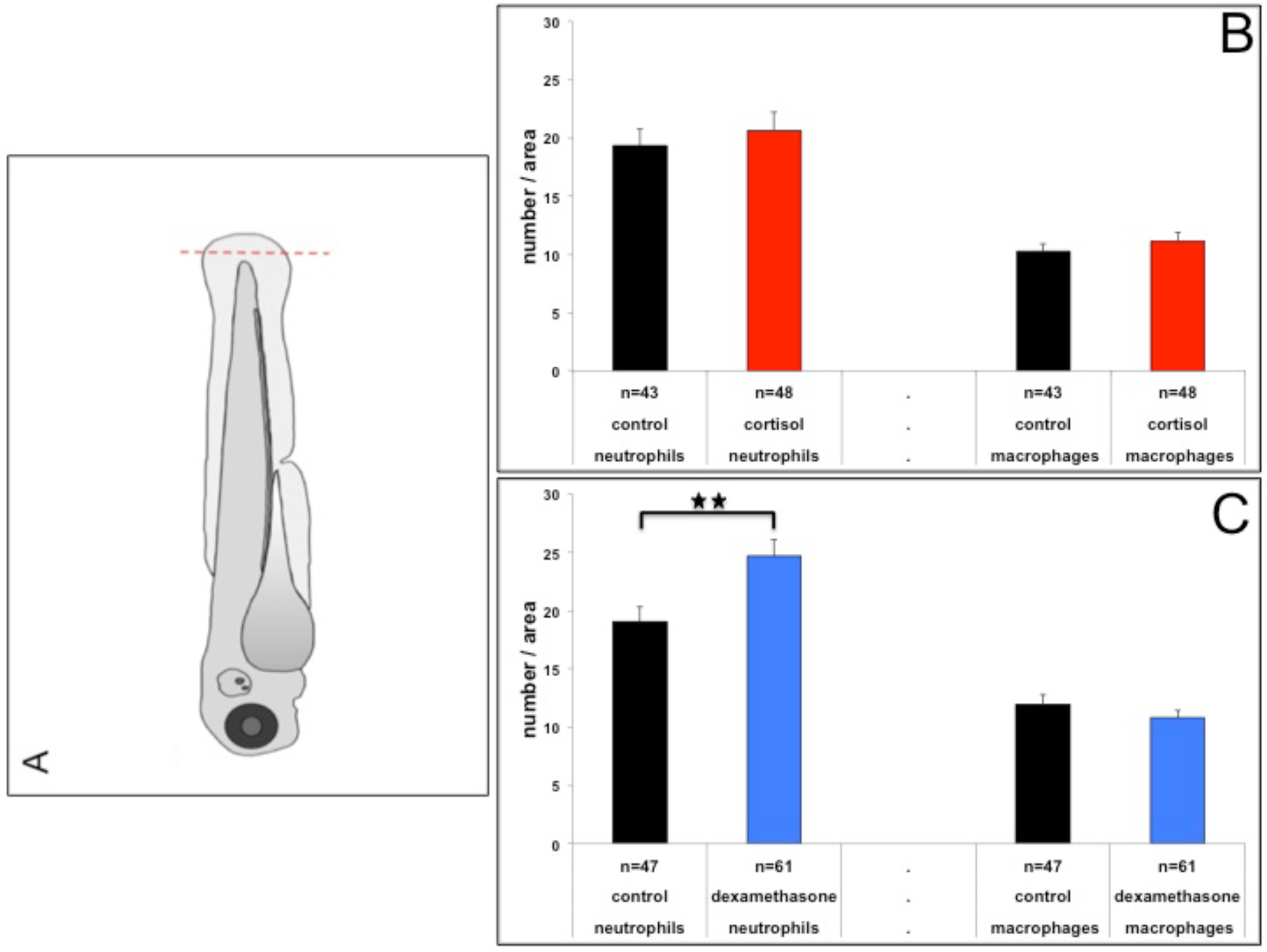
**panel A** Picture that illustrates the position of the site of the tail fin amputation in 3 dpf larvae. **panel B**: Number of neutrophils and macrophages (mean± SEM) at 4 hpa for the different treatments (cortisol or control). **panel C**: Number of neutrophils and macrophages (mean± SEM) at 4 hpa for the different treatments (dexamethasone or control).

### 3.3 LPS exposure

#### Survival

In the LPS exposure experiments, we never observed any morphological changes or mortality in the 4 dpf control groups. Hence, we only present the data of the 4 dpf LPS-treated groups. Figure 3 shows the survival data of both LPS exposure experiments. Only 27.8% of the 0-6 hpf control-treated larvae survived in the first LPS exposure experiment (figure 4A), while 79.2% survived in the second LPS exposure experiment (figure 4B). In both LPS exposure experiments the number of larvae that survived following glucocorticoid treatment 0-6 hpf appeared to be higher (Log Rank Mantel Cox; first LPS exposure experiment: overall Chi-square=20.863, df=2, p<0.001; second LPS exposure experiment: overall Chi-square=5.824, df=2, p≤0.054). Pair-wise comparison revealed that (1) in both LPS exposure experiments survival was significantly higher in 0-6 hpf cortisol-treated than 0-6 hpf control-treated larvae ((Log Rank Mantel Cox; Chi-square=14.385, df=1, p<0.001; Chi-square=5.466, df=1, p≤0.019), while (2) survival was significantly higher in the first LPS exposure experiment (Log Rank Mantel Cox; Chi-square=12.436, df=1, p<0.001), but not the second LPS exposure experiment (Log Rank Mantel Cox; Chi-square=1.370, df=1, ns) in 0-6 hpf dexamethasone-treated larvae compared to 0-6 hpf control-treated larvae.

**Figure 4.**
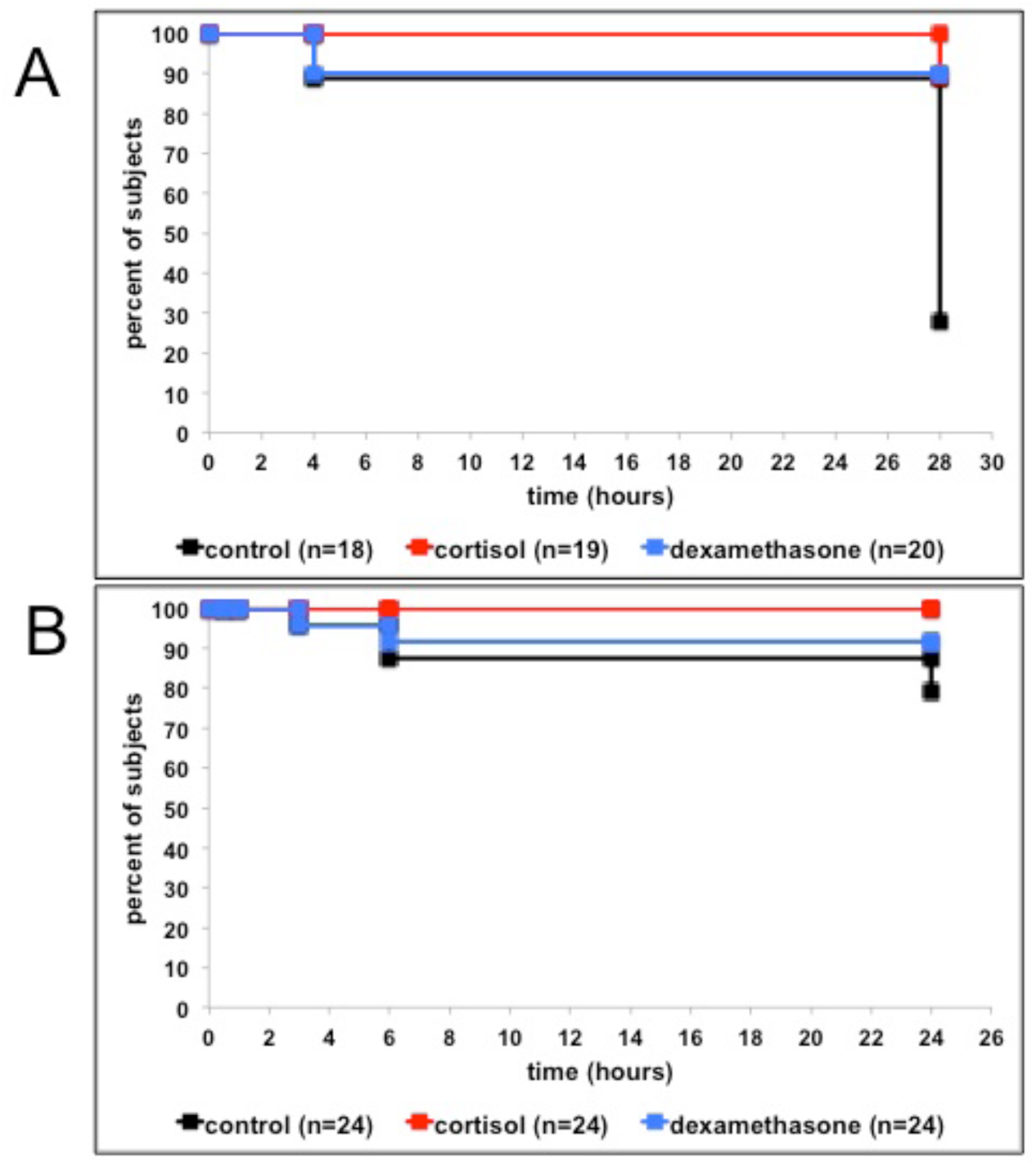
**panel A:** Per cent of surviving larvae following LPS treatment (from 0-0.5 hrs) and 0-6 hpf control treatment, cortisol treatment or dexamethasone treatment in the first exposure series (time-points: 0 hr, 4 hrs and 28 hrs). **panel B**: Per cent of surviving larvae following LPS treatment (from 0-0.5 hrs) and 0-6 hpf control treatment, cortisol treatment and dexamethasone treatment in second exposure series (time-points: 0hr, 0.5 hr, 1 hr, 3 hrs, 6 hrs and 24 hrs).

#### Phenotypical effects and gene expression endothelium-related genes

In the first LPS exposure experiment phenotypical changes were measured at 4 and 28 hrs, i.e. 3.5 hrs and 27.5 hrs after ending exposure. Table 2 shows the numbers of dead larvae and of larvae that were alive and displayed phenotypical changes. We observed LPS-induced changes in the shape of the larvae (curved larvae), tail fin morphology (swollen or damaged tail fins) and heart cavity (oedema). Larvae were scored as either affected (at least one of these changes present) or not (no changes in any of the parameters). Supplementary table 1 shows the scores of the individual parameters at 4 hrs (the number of larvae of the control treated group alive at 28 hrs was too low for a meaningful statistical analysis between treatments).

**Table 2:** Per cent of larvae dead, alive and affected (regardless of phenotypical changes in the tail fin, shape or cardiac area), and alive and intact at 4hrs and 28hrs after LPS treatment (0-0.5hr) in the different treatment groups: 0-6 hpf treatment with control, cortisol-containing or dexamethasone-containing medium. Note that the number of larvae at 28hrs is lower than at 4hrs; per treatment n=3-4 larvae (alive) were randomly selected for staining for reactive oxygen species in the tail fin (not reported here); they were not scored for phenotypical changes.

| 4 hrs | treatment | %dead | %(alive+affected)<br>(tail fin, shape, heart) | %(alive+intact) |
| --- | --- | --- | --- | --- |
|  | control (n=18) | 11.1 | 83.3 | 5.6 |
|  | cortisol (n=19) | 0.0 | 73.7 | 26.3 |
|  | dexamethasone (n=20) | 10.0 | 60.0 | 30.0 |
| 28 hrs | treatment | %dead | %(alive+affected)<br>(tail fin, shape, heart) | %(alive+intact) |
|  | control (n=15) | 86.7 | 13.3 | 0.0 |
|  | cortisol (n=16) | 12.5 | 37.5 | 50.0 |
|  | dexamethasone (n=16) | 12.5 | 31.3 | 56.3 |

While the scores at 4 hrs suggested that the LPS-induced effects were less strong in glucocorticoid-treated larvae than in control-treated larvae, this was not (as yet) significant (Chi-square=5.81, df=2, ns). LPS-induced effects were less strong in glucocorticoid-treated larvae than in control-treated larvae at 28 hrs (overall Chi-square=25.33, df=2, p<0.001). Pair-wise comparison showed that both cortisol-treated (Chi-square=18.05, df=2, p<0.001) and dexamethasone-treated (Chi-square=18.34, df=2, p<0.001) groups showed fewer dead and fewer malformed larvae than control-treated subjects following LPS exposure.

In addition to the phenotypical changes that we studied, we measured expression levels of genes related to endothelial barrier function (*clnd5a*, *clnd2* and *oclnb*). Transcript abundance of *clnd5a* was higher in cortisol-treated (Tukey HSD: p<0.001) and dexamethasone-treated larvae (Tukey HSD: p≤0.067) compared to control-treated larvae (two-way ANOVA; treatment and time as independent factor; treatment: F(2,36)=9.815, p<0.001; figure 5A). Transcript abundance of *clnd2* was lower in cortisol-treated (Tukey HSD: p≤0.034) and dexamethasone-treated larvae (Tukey HSD: p≤0.093) compared to control-treated larvae (two-way ANOVA; treatment and time as independent factor; treatment: F(2,36)=9.815, p<0.001; figure 5B). At 3 hrs transcript abundance of *clnd2* was higher than at other time points in all treatment groups (Tukey HSD; p<0.001; time: F(2,36)=13.425, p<0.001). Transcript abundance of *oclnb* was lower in cortisol-treated and dexamethasone-treated larvae than in control-treated larvae at baseline (0 hr), but higher at 3 hrs following LPS exposure (two-way ANOVA; treatment and time as independent factor; treatment *time: F(3,36)=6.404, p<0.001; figure 5C).

**Figure 5.**
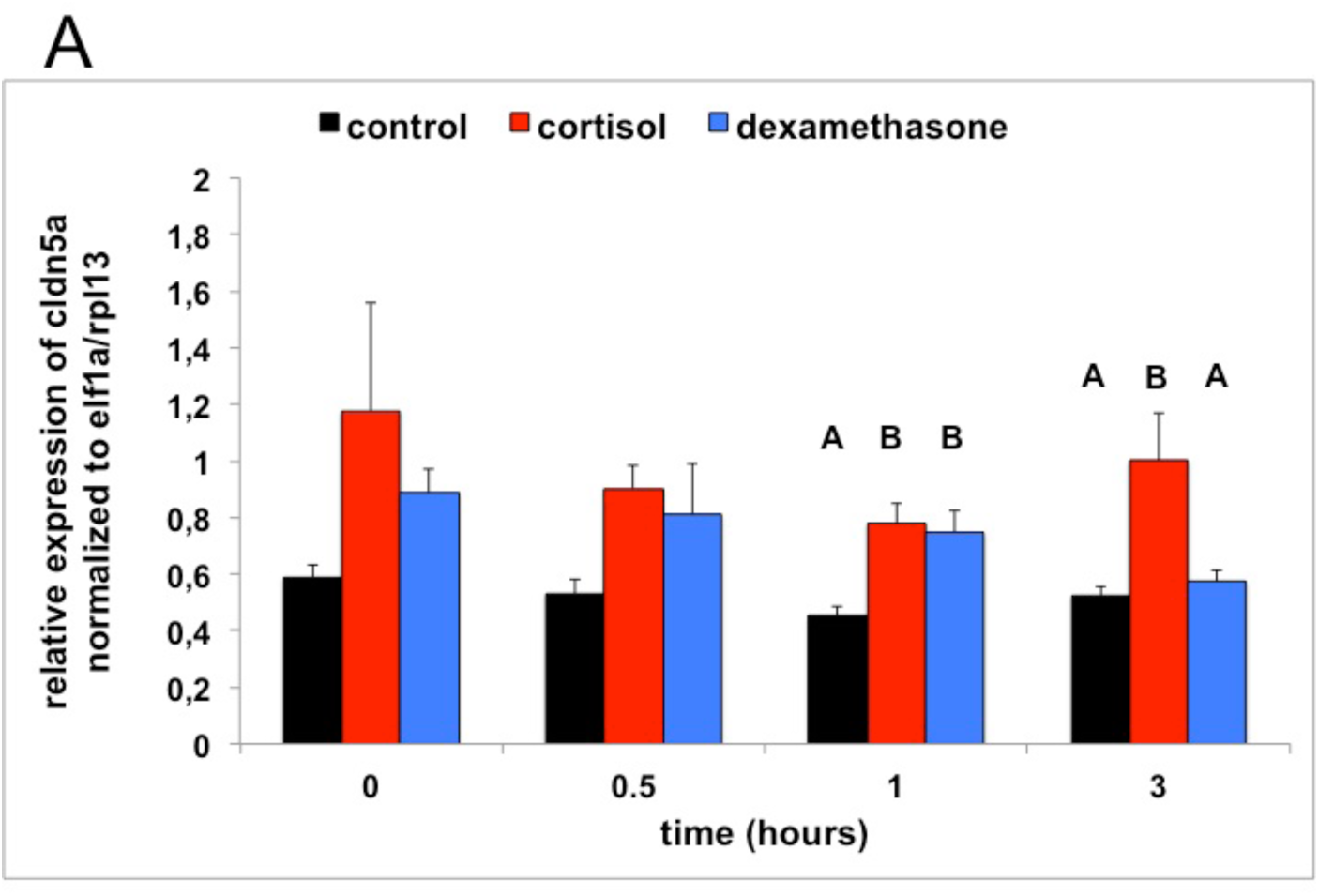
**panel A:** Transcript abundance (relative normalized expression; mean+SEM) of *cldn5a* following LPS treatment (from 0-0.5 hr) and 0-6 hpf control treatment, cortisol treatment or dexamethasone treatment (n=4 samples per time-point). Groups with the same capitals do not significantly differ from one another (Tukey HSD following a significant treatment effect for this time-point). Note: control-treated subjects showed no significant change over time: F(3,12)=2.009, ns.

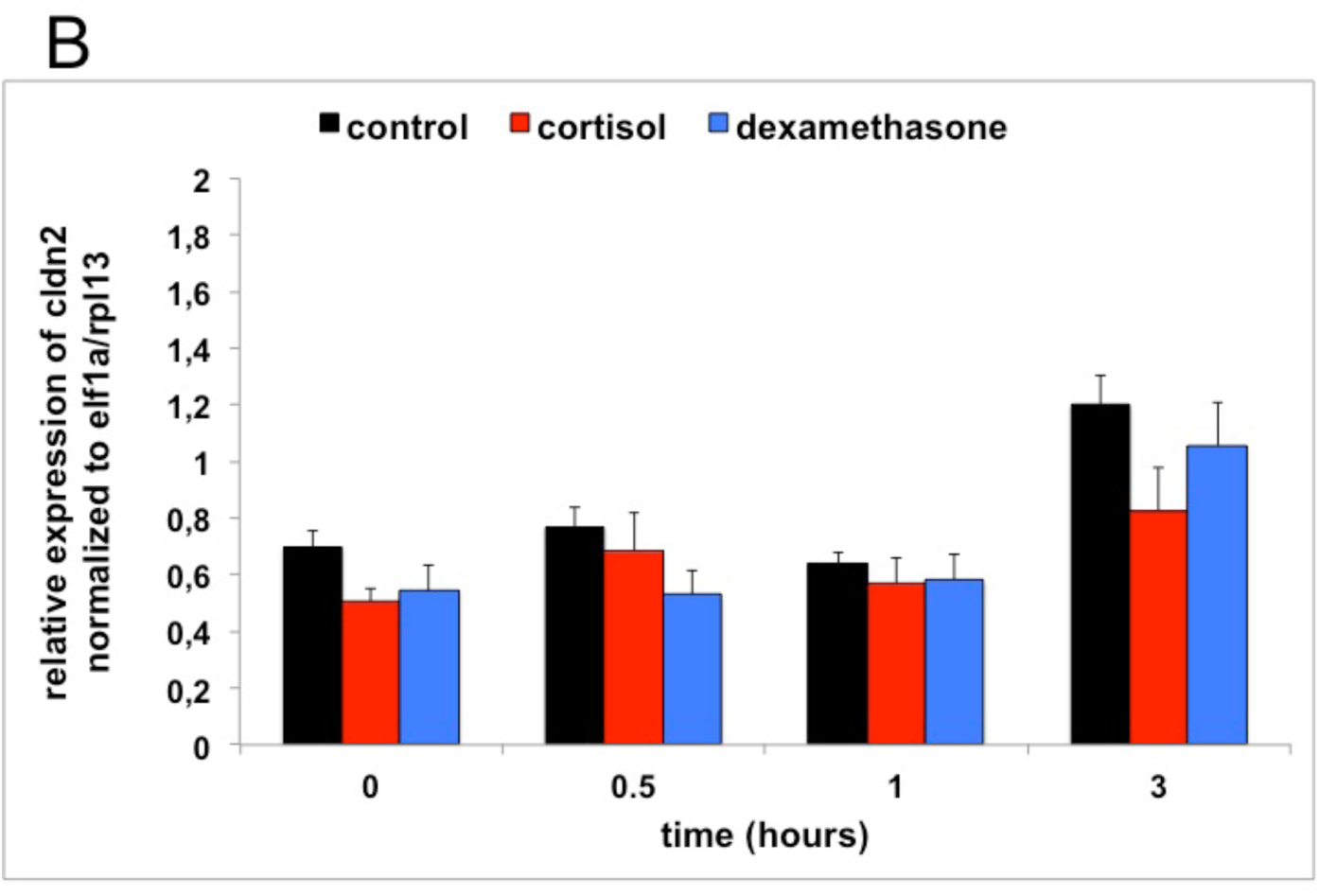
**panel B:** Transcript abundance (relative normalized expression; mean+SEM) of *cldn2* following LPS treatment (from 0-0.5 hr) and 0-6 hpf control treatment, cortisol treatment or dexamethasone treatment (n=4 samples per time-point). Note: control-treated subjects showed an increased expression at 3 hrs: F(3,12)=13.253, p<0.001; Tukey HSD.

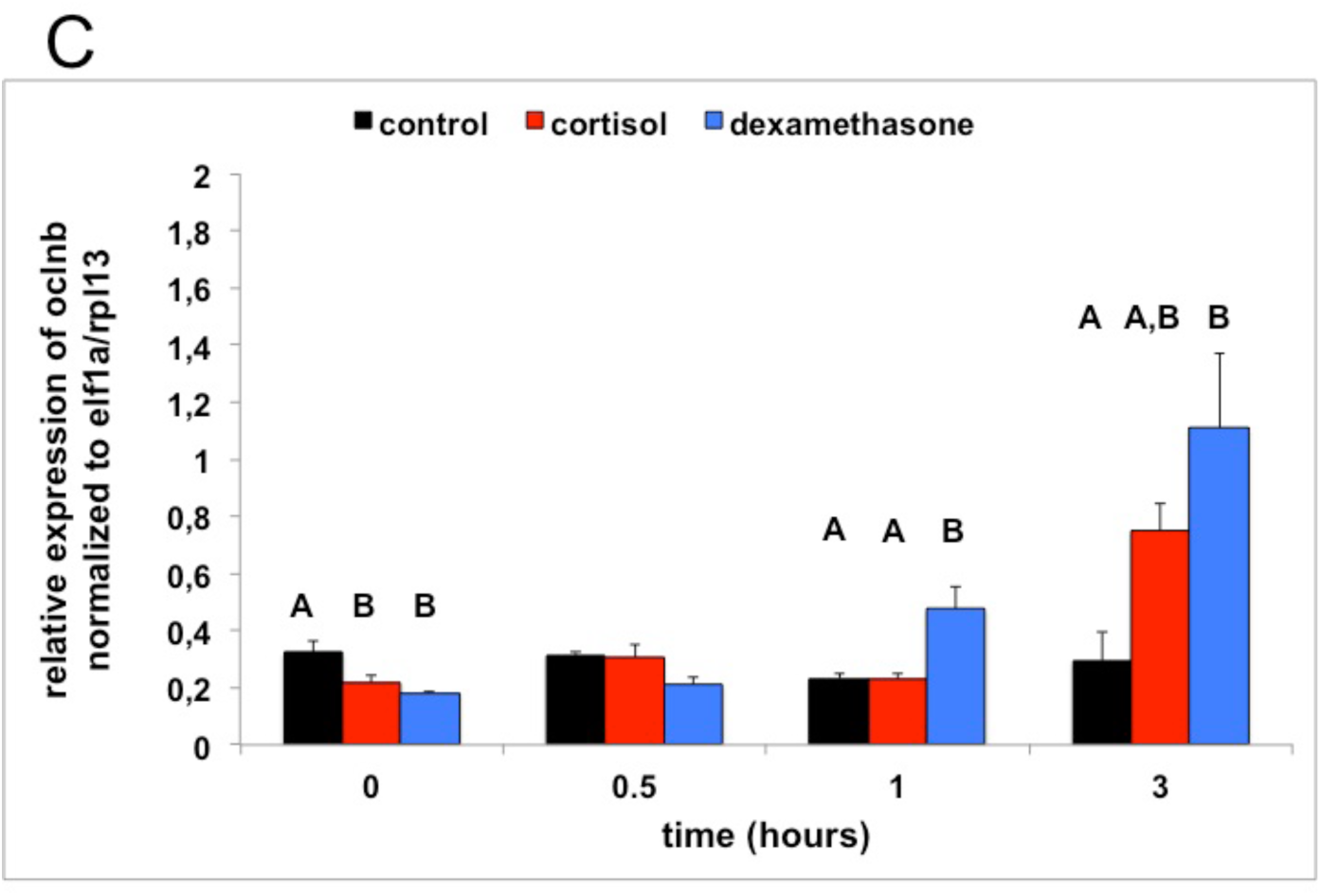
**panel C**: Transcript abundance (relative normalized expression; mean+SEM) of *oclnb* following LPS treatment (from 0-0.5 hr) and 0-6 hpf control treatment, cortisol treatment or dexamethasone treatment (n=4 samples per time-point). Groups with the same capitals do not significantly differ from one another (Tukey HSD following a significant treatment effect for this time-point). Note: control-treated subjects showed no significant change over time: F(3,12)=0.562, ns.

In the second LPS exposure experiment we measured changes in tail fin morphology and shape as we hardly observed any oedema in the heart cavity in the first LPS exposure experiment. Figure 6 shows the changes in tail fin morphology following LPS exposure. In control-treated larvae there was a clear and rapid loss of normal tail fin structure and a shift towards swollen or damaged tail fins, while this was not the case in the glucocorticoid-treated larvae. Statistical analysis showed that at 3 hrs (Chi-square=10.25, df=4, p≤0.036), 6 hrs (Chi-square=10.27, df=4, p≤0.036) and 24 hrs (Chi-square=10.25, df=4, p≤0.042) treatment groups differed significantly from one another. More in particular, both cortisol-treated larvae (3 hrs: Chi-square=5.16, df=2, p≤0.076; 6 hrs: Chi-square=6.94, df=2, p≤0.03; 24 hrs Chi-square=5.77, df=2, p≤0.056) and dexamethasone-treated larvae (6 hrs: Chi-square=6.98, df=2, p≤0.03; 24 hrs Chi-square=6.53, df=2, p≤0.038) showed less severe changes in tail fin morphology than control-treated larvae. We did not observe a strong effect of LPS exposure on the shape of the larvae. After 24 hrs only small percentages of each treatment group showed a curved shape: control-treated larvae (see supplementary table 2 for all time points): 15.8% (n=19); cortisol-treated larvae: 8.3% (n=24); dexamethasone-treated larvae: 9.1% (n=22); these differences were not significant (Chi-square=1.71, df=2, ns).

**Figure 6.**
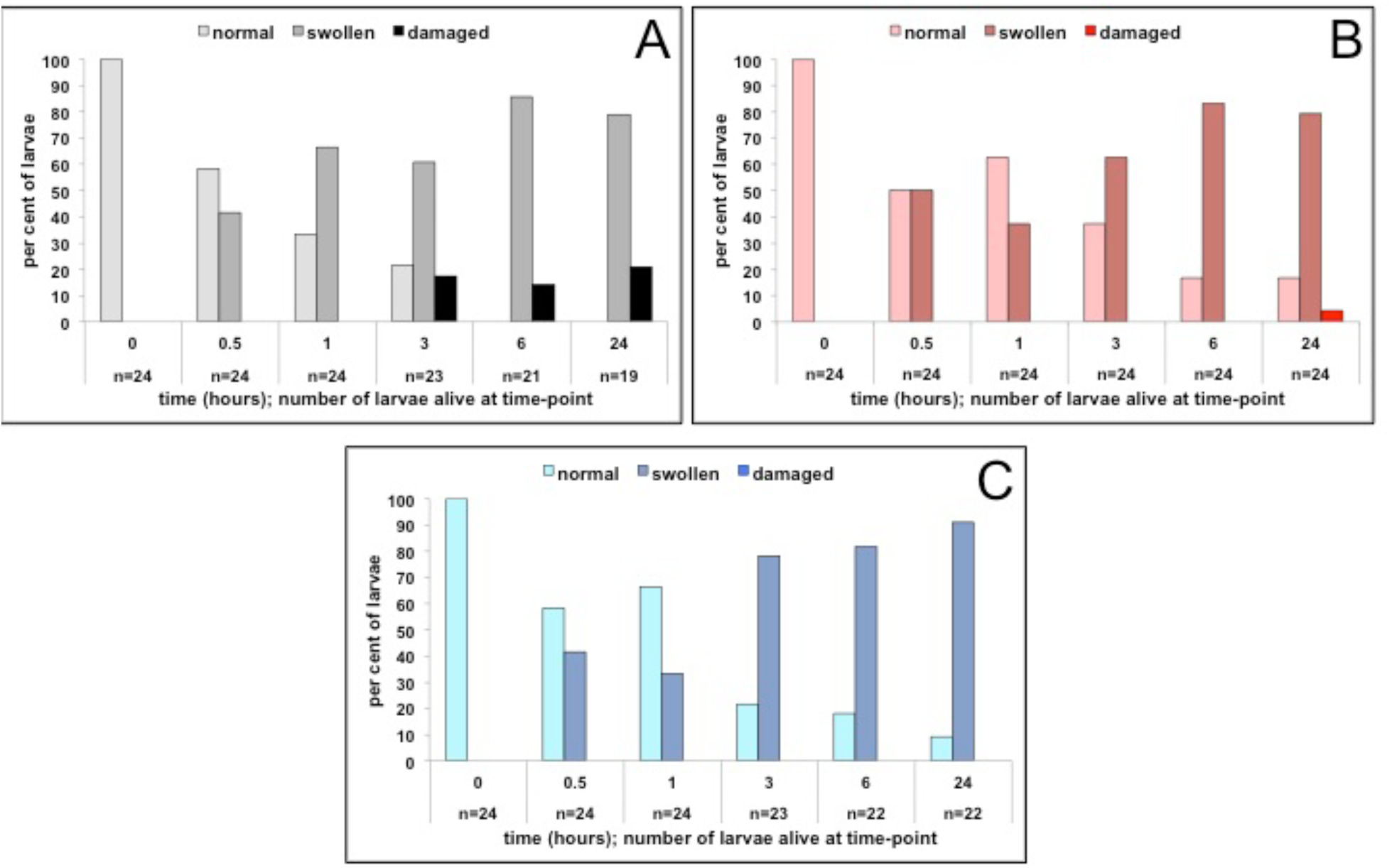
**panel A**: Per cent of larvae following LPS treatment (from 0-0.5 hr) in the 0-6 hpf control treatment group showing normal, swollen or damaged tail fins (time-points: 0hr, 0.5hr, 1 hr, 3 hrs, 6 hrs and 24 hrs). **panel B**: Per cent of larvae following LPS treatment (from 0-0.5 hr) in the 0-6 hpf cortisol treatment group showing normal, swollen or damaged tail fins (time-points: 0 hr, 0.5 hr, 1 hr, 3 hrs, 6 hrs and 24 hrs). **panel C**: Per cent of larvae following LPS treatment (from 0-0.5 hr) in the 0-6 hpf dexamethasone treatment group showing normal, swollen or damaged tail fins (time-points: 0 hr, 0.5 hr, 1 hr, 3 hrs, 6 hrs and 24 hrs).

#### Gene expression analysis

First, we explored the effects of glucocorticoid treatment through the interrelationships of transcript abundance of genes using PCA. Then in each component, we selected genes of interest of which the transcript abundance of the different treatments matched the differences in phenotypical changes and survival that we observed between treatment groups.

A PCA revealed three components explaining in total 72.9% of variance (supplementary table 3). The KMO was sufficiently high (0.619) and Bartlett’s test of sphericity was highly significant (Chi-square=303.010, df=55, p<0.001) indicating that the data were adequate for a PCA.

The first component, explaining 32.9% of the variance, consisted of the genes of different Toll-like receptors: *tlr2* (factor loading: 0.677), *tlr4ba* (0.669), *tlr4bb* (0.764), *tlr5a* (0.779) and *tlr5b* (0.838). The overall pattern of the factor regression scores across time was an inverted U-shape: a two-way ANOVA (independent factors: time and treatment) for this component revealed a highly significant effect of time (F(3,36)=15.502, p<0.001) with time points 0 hr and 3 hrs having significantly higher factor regression scores than time points 0.5 hr and 1 hr (Tukey HSD; supplementary Table 3). Only weak effects were found between the different treatments (F(2,36)=3.012, p≤0.062; F(6,36)=2.038, p≤0.086). The only gene of this component of which the transcript abundance of the different treatments seemed to match the differences in phenotypical changes and survival was *tlr4bb* (see supplementary table 4 and figure 7A): transcript abundance increased less strongly over time in glucocorticoid-medium treated subjects than in control-medium treated subjects.

**Figure 7.**
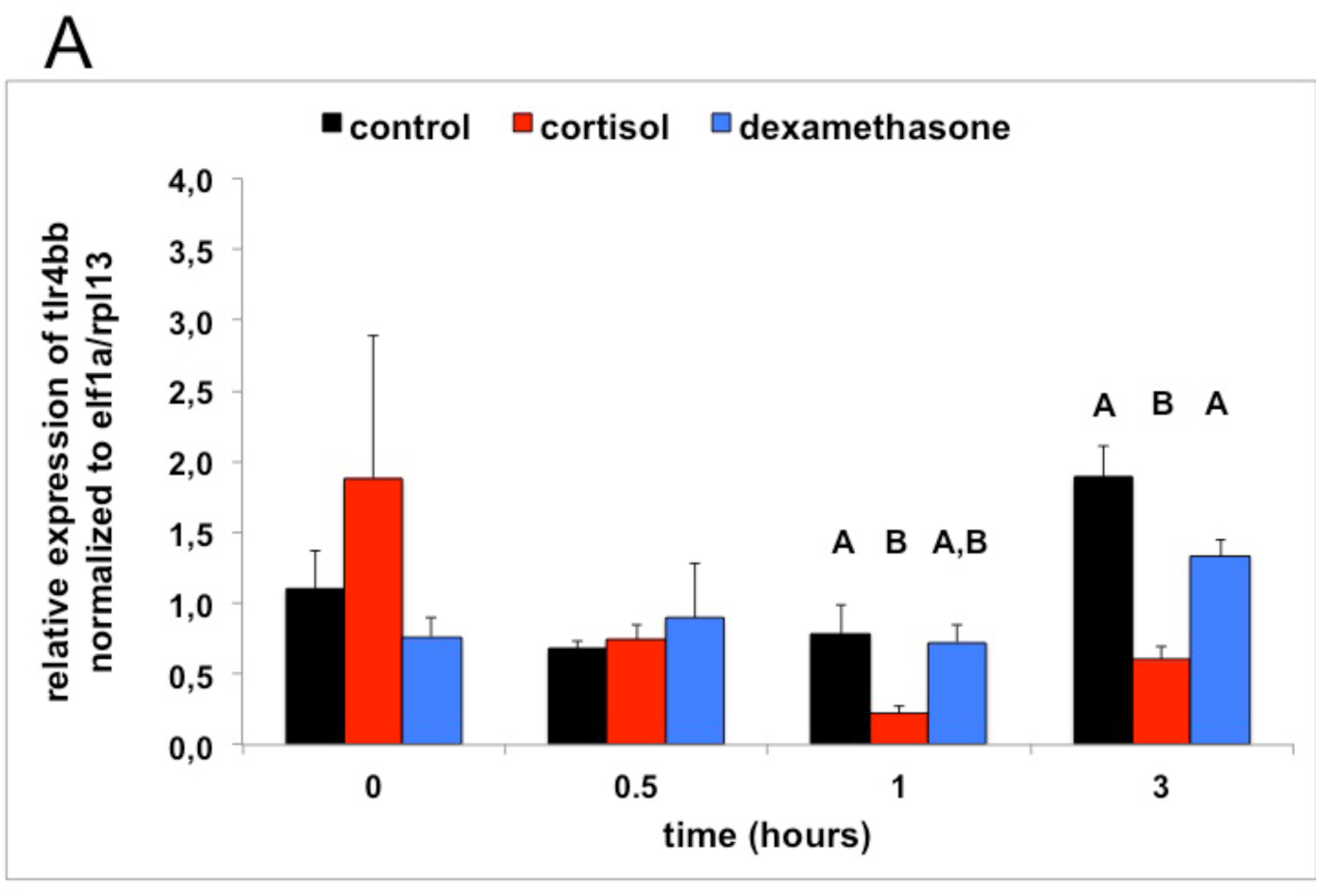
**panel A**: Transcript abundance (relative normalized expression; mean+SEM) of *tlr44bb* following LPS treatment (from 0-0.5 hr) and 0-6 hpf control treatment, cortisol treatment or dexamethasone treatment (n=4 samples per time-point). Groups with the same capitals do not significantly differ from one another (Tukey HSD following a significant treatment effect for this time-point).

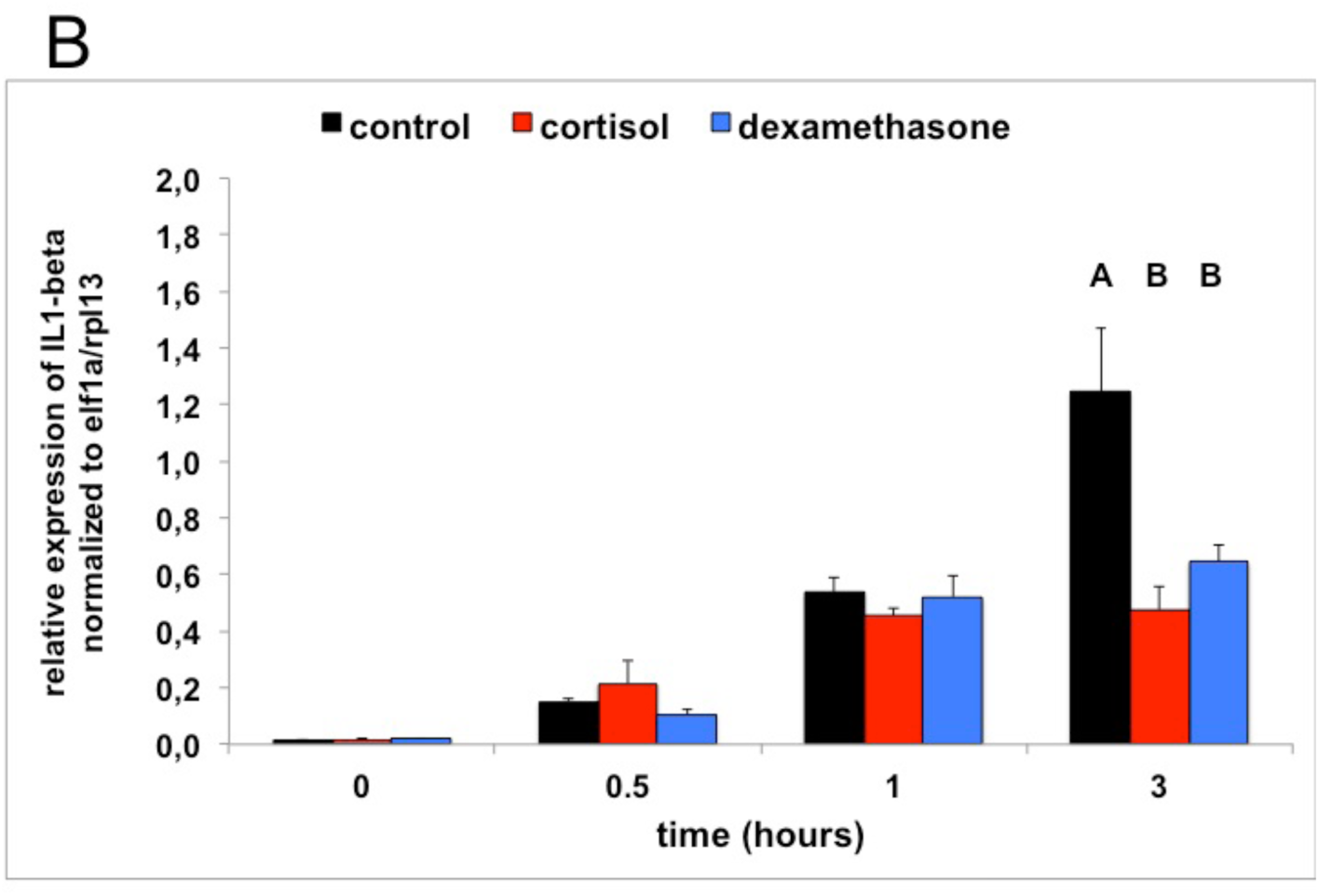
**panel B**: Transcript abundance (relative normalized expression; mean+SEM) of *IL1-beta* following LPS treatment (from 0-0.5 hr) and 0-6 hpf control treatment, cortisol treatment or dexamethasone treatment (n=4 samples per time-point). Groups with the same capitals do not significantly differ from one another (Tukey HSD following a significant treatment effect for this time-point).

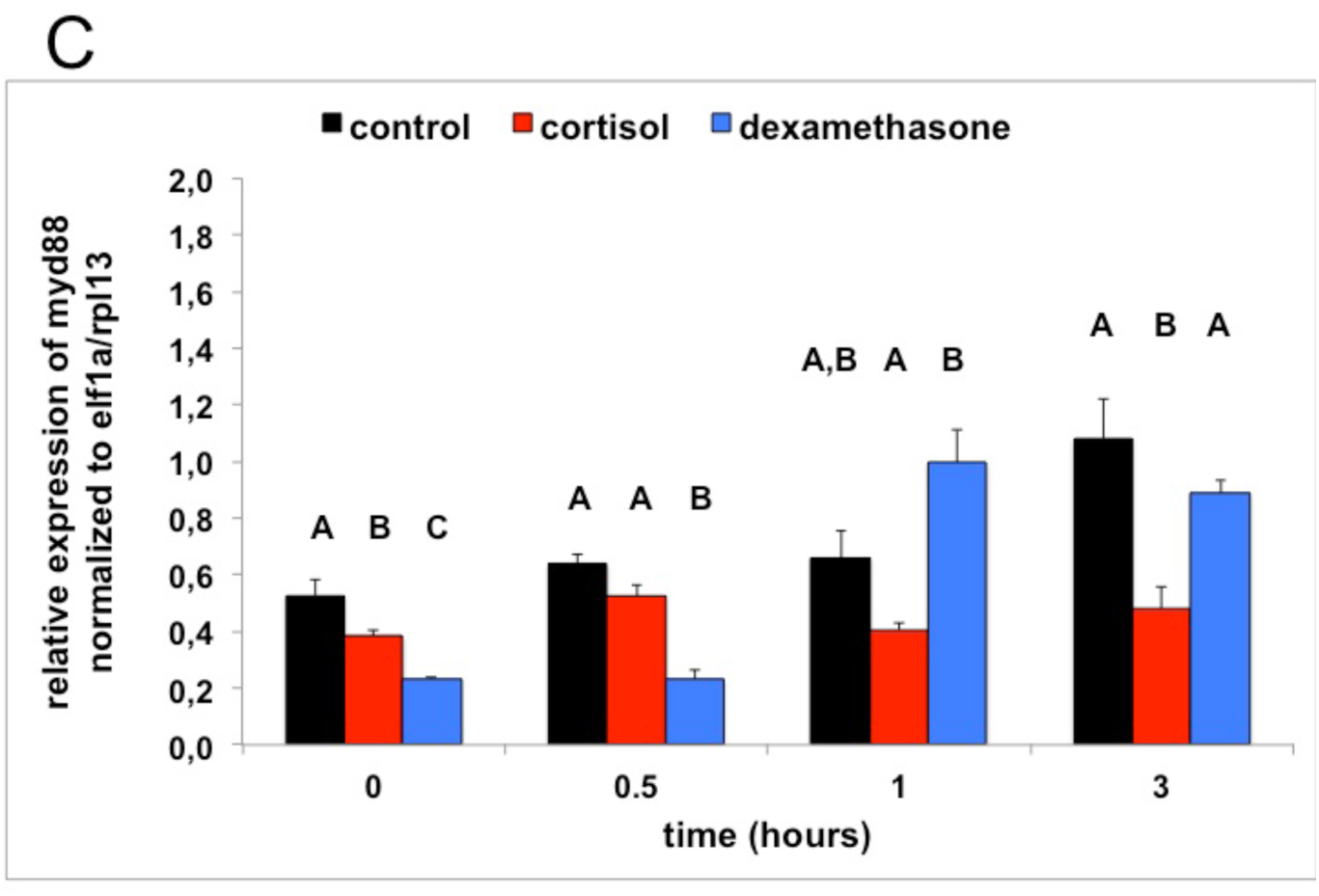
**panel C**: Transcript abundance (relative normalized expression; mean+SEM) of *myd88* following LPS treatment (from 0-0.5 hr) and 0-6 hpf control treatment, cortisol treatment or dexamethasone treatment (n=4 samples per time-point). Groups with the same capitals do not significantly differ from one another (Tukey HSD following a significant treatment effect for this time-point).

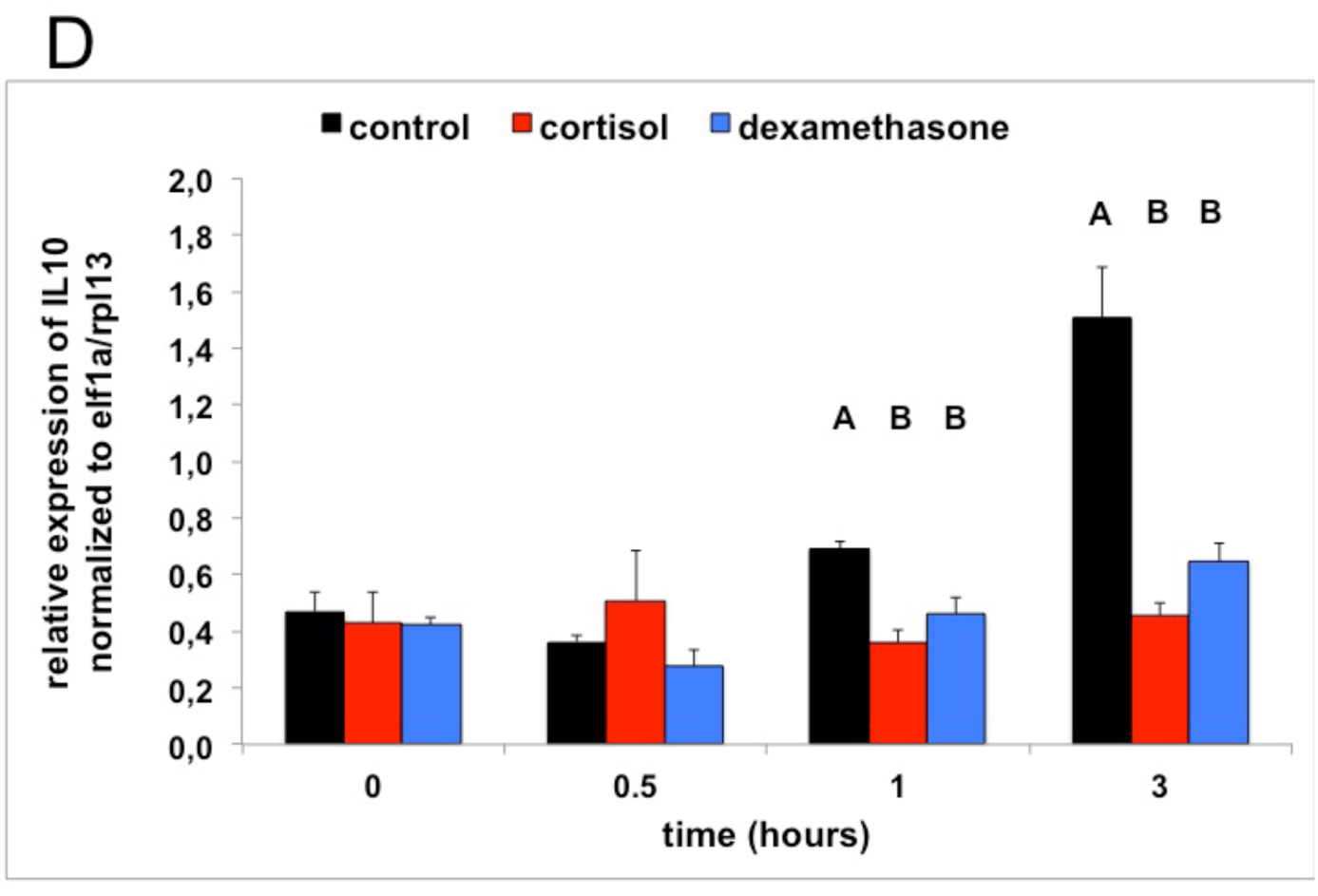
**panel D**: Transcript abundance (relative normalized expression; mean+SEM) of *IL10* following LPS treatment (from 0-0.5 hr) and 0-6 hpf control treatment, cortisol treatment or dexamethasone treatment (n=4 samples per time-point). Groups with the same capitals do not significantly differ from one another (Tukey HSD following a significant treatment effect for this time-point).

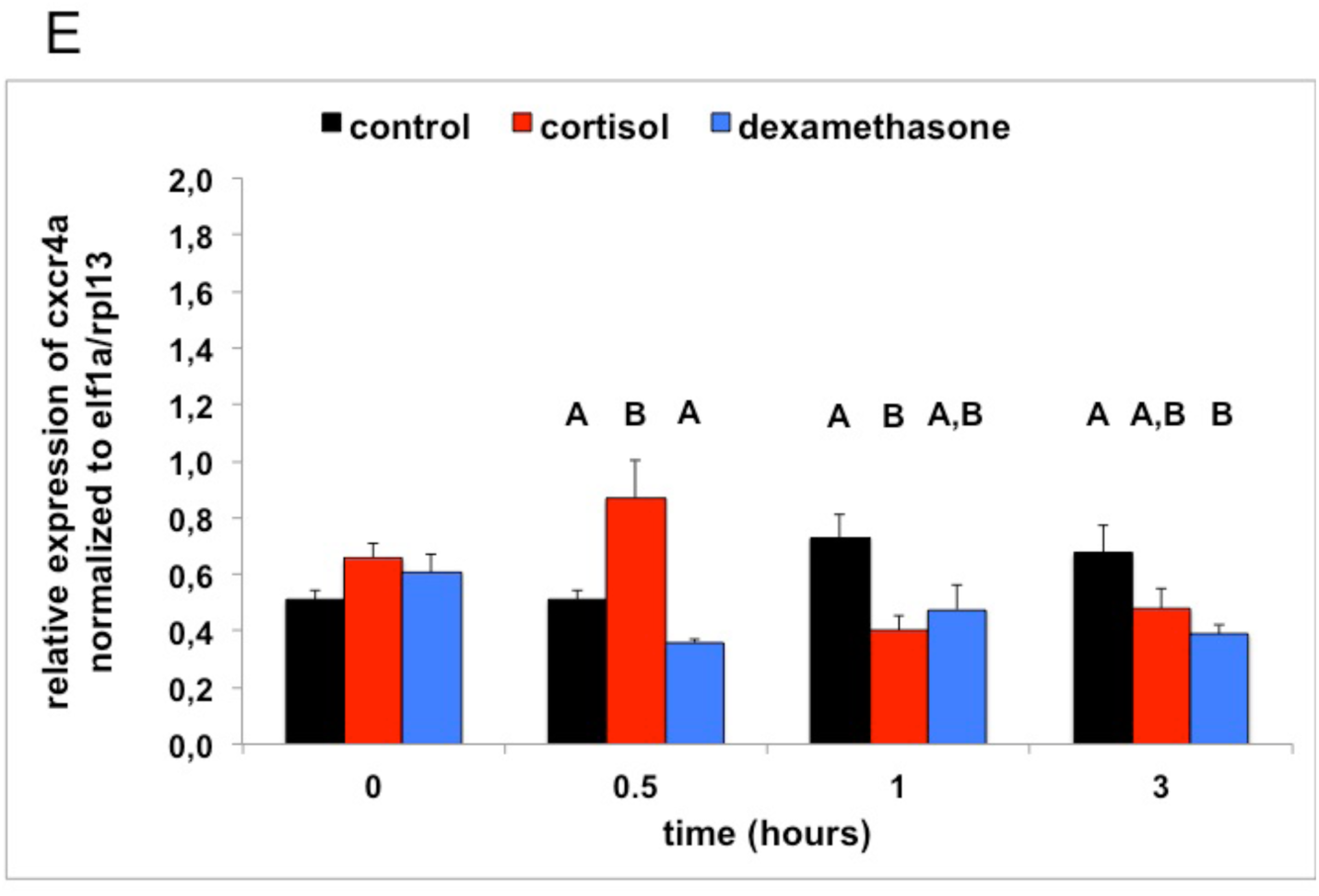
**panel E**: Transcript abundance (relative normalized expression; mean+SEM) of *cxcr4a* following LPS treatment (from 0-0.5 hr) and 0-6 hpf control treatment, cortisol treatment or dexamethasone treatment (n=4 samples per time-point). Groups with the same capitals do not significantly differ from one another (Tukey HSD following a significant treatment effect for this time-point).

The second component, explaining 23.1% of the variance, consisted of *il1β* (factor loading: 0.940), *myd88* (0.844) and *il10* (0.758). As time progressed factor regression scores increased (two-way ANOVA (independent factors: treatment and time); time: F(3,36)=68.309, p<0.001) with a post-hoc Tukey HSD revealing that the factor regression scores differed significantly from one another at all time points (supplementary table 3). The 0-6 hpf control group had increasingly higher factor regression scores than the 0-6 hpf cortisol treated group and the 0-6 hpf dexamethasone treated group as time progressed (supplementary table 3; Tukey HSD; treatment: F(2,63)=22.664, p<0.001; treatment*time: F(6,36)=13.491, p<0.001). Of all genes of this component, transcript abundance of the different treatments seemed to match the differences in phenotypical changes and survival (supplementary table 4 and figure 7 B–D): transcript abundance increased less strongly over time in glucocorticoid-medium treated subjects than in control-medium treated subjects.

The third component, explaining 16.9% of the variance, consisted of regulators of the immune response: *cxcr4a* (factor loading: 0.794), *cxcr4b* (0.809) and *ptpn6* (0.799). Overall the factor regression scores decreased over time: a two-way ANOVA (independent factors: treatment and time) revealed a highly significant effect of time (F(3,36)=4.464, p≤0.009) with 3 hrs and 0 hr being significantly different from one another (Tukey HSD). The 0-6 hpf dexamethasone group had lower factor regression scores than the 0-6 hpf control treated group and the 0-6 hpf cortisol treated group (treatment: F(2,36)=11.056, p<0.001; treatment*time: F(6,36)=5.571, p<0.001). The only gene of this component of which the transcript abundance of the different treatments seemed to match the differences in phenotypical changes and survival was *cxcr4a* (see supplementary table 4 and figure 7E): transcript abundance increased in control-medium treated subjects but decreased in glucocorticoid-medium treated subjects.

## 4. Discussion

The data of this study showed that treatment of zebrafish embryos with cortisol or dexamethasone during the first six hours after fertilization modulated the function of the immune system and thereby enhanced survival after an immune challenge. This suggests that in zebrafish maternal stress through enhancing oocyte cortisol levels and thereby increased GR stimulation leads to an adaptive response to immune challenges.

### 4.1. Baseline expression of immune-related genes

Hartig and colleagues (2016) have shown that 5-day exposure to (1 µM) cortisol in zebrafish embryos/larvae (0-5 dpf) enhanced baseline expression of immune-related genes such as of *socs3a*, *mpeg1* and *irg1l* at 5 dpf. Here, we show that exposure at the first six hours of life (0-6 hpf) is already sufficient to induce enhanced baseline expression of these genes at 5 dpf. As GR is the only corticosteroid receptor present in these early life stages (Alsop and Vijayan, 2008; Pikulkaew *et al*., 2010, 2011), this suggests that activation of GR in these early stages is responsible for mediating these effects. Indeed, we show that 0-6 hpf exposure to the specific GR agonist dexamethasone enhanced baseline expression of these immune-related genes in 5 dpf larvae as well. This is in general agreement with data from studies showing that exposure of embryos/larvae to other GR agonists (for variable time-periods from fertilisation) increased baseline expression of (some of) these genes (Willi *et al*., 2018, 2019; Zhao *et al*., 2016).

We have previously observed that 0-6 hpf exposure to cortisol enhanced baseline levels of cortisol at 5 dpf in larvae of the AB strain but not in larvae of the TL strain (van den Bos *et al*., 2019). In addition, we have observed that 0-6 hpf exposure to dexamethasone had no effect on baseline levels of cortisol at 5 dpf in larvae of the AB or TL strain (Althuizen, 2018). Overall these data suggest that the effects on the expression of immune-related genes in AB and TL larvae are independent of baseline levels of cortisol.

While we have previously observed substantial differences between larvae of the AB and TL strains in gene expression, physiology and behaviour (van den Bos *et al*., 2017a, 2017b, 2019, 2020), the effects of 0-6 hpf exposure to cortisol and dexamethasone on baseline expression of immune-related genes were similar in both strains. This suggests a robust strain-independent effect. Future studies should elucidate whether this effect also holds in other strains revealing thereby fundamental aspects of how early-life levels of cortisol may affect offspring functioning and survival.

### 4.2 Tail fin amputation assay

It has been shown that tail fin amputation in 3 dpf larvae leads to a rapid recruitment of macrophages (within 2 hpa) remaining at a plateau for at least 24 hours thereafter, while the number of neutrophils reaches a peak 4 hpa declining thereafter (Chatzopoulou *et al*., 2016). Here, we observed that dexamethasone increased the number of neutrophils at 4 hpa, while it had no effect on the number of macrophages at 4 hpa. These data suggest that pre-exposed subjects mount a stronger response to injury, possibly enhancing protection against invading micro-organisms (Hall *et al*., 2014).

Dexamethasone, but not cortisol, exposure was effective in increasing the recruitment of neutrophils. Two explanations may be forwarded for this. First, this may be related to dexamethasone’s higher affinity for GR (Rupprecht *et al*., 1993). In line with this we observed enhanced expression of FK506 binding protein 5 (*fkbp5*), a marker for levels of GR stimulation (e.g. Willi *et al.,* 2018, 2019) in dexamethasone-exposed embryos, but not cortisol-exposed embryos (Althuizen, 2018; van den Bos *et al*., 2019). Second, it has been shown neutrophil recruitment, but not macrophage recruitment, is sensitive to acute treatment with GR agonists, including cortisol, which decreases the number of neutrophils at 4 hpa (Chatzopoulou *et al*., 2016; Hall *et al*., 2014; Xie *et al*., 2019). We have observed that baseline levels of cortisol are enhanced following cortisol but not dexamethasone treatment (Althuizen, 2018; van den Bos *et al*., 2019). Hence, in the case of cortisol pre-exposure enhancement of neutrophil recruitment through GR stimulation (as indicated by the effect of dexamethasone) may potentially be offset at 3 dpf by the higher baseline levels of cortisol. If so, this suggests a fine-tuning of the response. Future studies should address these alternative explanations.

### 4.3 LPS challenge

In line with results from other studies (Dios *et al*., 2014; Hsu *et al*., 2018; Novoa *et al*., 2009; Philip *et al*., 2017) LPS exposure produced phenotypical changes (tail fin oedema, swollen or damaged tail fins, and curved animals), mortality and increased expression levels of immune-related genes. To assess the molecular mechanisms underlying the effects of LPS exposure, we measured the expression of *cldn5a*, *cldn2* and *oclnb*; genes of which the expression has been shown to be changed following LPS exposure (Hsu *et al*., 2018; Philip *et al*., 2017) and which are involved in endothelial barrier function (Kása *et al*., 2015; Odenwald and Turner, 2013; Shen *et al*., 2011; Yoseph *et al*., 2016). The strong increase in the expression of *cldn2* at 3 hours in control-treated subjects is in line with data from other studies (Hsu *et al*., 2018; Philip *et al*., 2017). Increased expression of *cldn2* is associated with endothelial hyper-permeability due to increased pore-pathway activity possibly mediated by increased expression of IL-13 (Kása *et al*., 2015; Odenwald and Turner, 2013; Shen *et al*., 2011; Yoseph *et al*., 2016). At variance with other studies (Hsu *et al*., 2018; Philip *et al*., 2017) we did not observe strongly decreased expressions of *cldn5a* and *oclnb*. Decreased expression levels of *cldn5a* and *oclnb* are related to endothelial hyper-permeability due a lower sealing function of the pore-pathway and a less functional leaky pathway respectively (Kása *et al*., 2015; Odenwald and Turner, 2013; Shen *et al*., 2011; Yoseph *et al*., 2016). One reason for this may be that we measured gene expressions at 3 hours after the start of the LPS exposure, while in other studies this was measured at substantially later time points, i.e. 6 and 8 hours (Hsu *et al*., 2018; Philip *et al*., 2017).

Both cortisol and dexamethasone exposure at 0-6 hpf were associated with milder effects to LPS exposure (in both experiments), as indicated by milder phenotypical changes (lower number of larvae expressing tail fin oedema, damaged tail fins or curved animals), lower mortality and lower gene expression of immune-related genes, such as *il1β*. As to the expression of endothelium-related genes we noted that the expression of *cldn5a* was higher and expression of *cldn2* lower in cortisol- and dexamethasone-treated subjects compared to control-treated subjects. This suggests a lower permeability of the endothelium due to a lower pore-pathway activity (Kása *et al*., 2015; Odenwald and Turner, 2013; Shen *et al*., 2011; Yoseph *et al*., 2016) supporting the milder phenotypical effects following LPS exposure. The expression of *oclnb* was strongly increased after three hours in 0-6 hpf cortisol-treated and dexamethasone-treated subjects compared to control-treated subjects. This suggests lower permeability due to more protective leaky pathway activity (Kása *et al*., 2015; Odenwald and Turner, 2013; Shen *et al*., 2011; Yoseph *et al*., 2016). It has been shown that 24 hours following LPS challenge in zebrafish larvae the expression of *oclnb* is strongly up-regulated facilitating tissue-repair (Hsu *et al*., 2018). Again, the data support the milder phenotypical effects that we see following LPS exposure in cortisol-treated and dexamethasone-treated subjects compared to control-treated subjects.

To explore the underlying mechanisms of glucocorticoid treatment we measured the expression of a series of genes of interest and related their expression to the outcome of the phenotypical changes and mortality. Five genes stood out in this respect, reflecting two receptors (*tlr4bb* and *cxcr4a)*, a factor involved in the transduction pathway of the expression of cytokines (*myd88*) and two cytokines (*il1β* and *il10*).

While LPS exerts its effects through transduction mechanisms following binding to TLR4 in mammals (Goulopoulou *et al*., 2015; Kása *et al*., 2015), in zebrafish this is not clear as yet: *tlr4ba* and *tlr4bb* have been suggested to be paralogues rather than homologues and TLR4BA and TLR4BB have thus far not been shown to be activated by LPS possibly by lack of a binding site for LPS (Sepulcre *et al*., 2009; Sullivan *et al*., 2009). However TLR4BB has been shown to be involved in inflammatory processes as *tlr4bb* transcript abundance is increased following tail fin amputation (Chatzopoulou *et al*., 2016). Here, we observed an increase in *tlr4bb* expression in 0-6 hpf control-treated subjects over time, accompanied by a strong inflammatory response, which was less strong in 0-6 hpf cortisol-treated or dexamethasone-treated subjects, accompanied by a milder inflammatory response. This suggests a role for TLR4BB in the LPS-induced response.

In humans CXCR4 has been implicated in recognition of LPS or being part of a ‘LPS sensing apparatus’ in addition to TLR4 (Triantafilou *et al*., 2001, 2008). Furthermore, LPS increases the expression of *cxcr4* through an NF-κB signalling pathway associated with increased micro-vascular leakage in the lungs (Konrad *et al*., 2017) or increased colorectal tumor metastasis (Liu *et al*., 2017). Here, we observed an increase in *cxcr4a* expression in 0-6 hpf control-treated subjects over time, which was absent in 0-6 hpf cortisol-treated or dexamethasone-treated subjects. This difference in gene expression may be associated with differences in the extent of vascular leakage as suggested by the differences in the expression of genes involved in the endothelial barrier and differences in tail fin oedema between treatments as discussed above. In zebrafish *cxcr4a* is found in endothelial cells (blood vessels), while *cxcr4b* is not (Wei Chong *et al*., 2001), which may explain that we only observed a phenotype-related effect for the expression profile of *cxcr4a*. Interestingly, CXCR4 has been implicated in the development of tolerance to lethal doses of LPS in zebrafish larvae (Dios *et al*., 2014; Novoa *et al*., 2009). Thus, this suggests that CXCR4 is involved in modulating the response to LPS.

Overall therefore our data warrant further studies into the role of TLR4BB and CXCR4 in LPS-induced sepsis in zebrafish as well as into the effects of early life glucocorticoid stimulation hereon.

Earlier studies have shown that MYD88 knockout larvae show enhanced survival to LPS challenge (Hsu *et al*., 2018) and no increase in *il1β* expression (van der Vaart *et al*., 2013). Lethal, but not sub-lethal, doses of LPS have been found to be associated with high expression levels of *il1β* and *il10* in zebrafish larvae, indicative of a hyper-inflammatory response (Dios *et al*., 2014). In line with these findings we observed that the expression levels of *myd88*, *il1β* and *il10* were strongly increased 3 hours after LPS challenge associated with low survival in 0-6 hpf control-treated subjects, but lower levels of expression of all three genes with higher survival in 0-6 hpf cortisol-treated and dexamethasone-treated subjects. Myd88 is an adaptor protein critical to toll-like receptor signalling (except for TLR3; Goulopoulou *et al*., 2016) and IL1β receptor signalling (see Kanwal *et al*., 2013; van der Vaart *et al*., 2013) and thereby cytokine expression. It should be noted that *myd88* expression was already low in 0-6 hpf cortisol-treated and dexamethasone-treated subjects, suggesting lower transduction pathway activity, possibly leading to a lower stimulation of inflammatory pathways. It is clear that this deserves further study.

As indicated above following cortisol or dexamethasone treatment at 0-6 hpf we noted increased base-line expression levels of *irg1l*, *socs3a*, *mpeg1.1* and *mpeg1.2* compared to control treatment at 0-6 hpf. These increased base-line levels may aid in increased clearance of bacteria and preventing excessive inflammation and hence aid in increasing survival (Benard *et al*., 2015; Hall *et al*., 2014; Jo *et al*., 2005).

The data of LPS exposure in larvae observed here seem to match the data of LPS exposure in adult zebrafish following 0-5 dpf exposure to cortisol: LPS exposure did not increase *il1β* expression in different tissues measured (Hartig *et al*., 2016). Unfortunately no survival was measured in the latter study.

Overall these data suggest that the response to LPS of subjects pre-exposed to cortisol or dexamethasone is less strong than the response of subjects of pre-exposed to control medium. How differences in survival, phenotype and gene-expression levels are causally related remains to be studied. As we have observed that 0-6 hpf exposure to dexamethasone had no effect on baseline levels of cortisol at 5 dpf in larvae of the AB strain (Althuizen, 2018), the data suggest that these effects are independent of baseline levels of cortisol.

### 4.4 Limitations

A clear limitation is that we only used one dose of cortisol and dexamethasone. For convenience we used equimolar doses of cortisol and dexamethasone, which may have led to different levels of stimulation of GR 0-6 hpf. Thus, in future studies different dose-ranges may be warranted, e.g. to study whether higher concentrations of cortisol in the tail fin amputation assay have an effect on neutrophil recruitment.

Regarding the tail fin amputation assay it has been shown that glucocorticoids may play a role in the differentiation of macrophages into a pro-inflammatory (M1) phenotype (Xie *et al*., 2019). So, future studies should study in more detail the effects of early life exposure of glucocorticoids on the inflammatory response and wound healing. Similarly, we used LPS to induce a hyper-inflammatory response, i.e. sepsis (Hsu *et al*., 2018; Philip *et al*., 2017), as a model to study the effectiveness of our early life treatments. To assess the ecological relevance of our findings and their more general nature the effects of early-life exposure on larval exposure to different pathogens, such as of bacterial, viral or fungal origin, should be studied (see e.g. Meijer and Spaink, 2011; van der Vaart *et al*., 2013).

We observed a variable response to the LPS challenge in the two exposure series. This is not uncommon as we also observed variable responses in other immune-related paradigms such as the response to a dextran sodium sulphate (DSS) challenge (van den Bos *et al*., unpublished observations). While as yet speculative differences in baseline levels of expression of *il1β* may be one associated factor as we observed that higher baseline levels seem associated with milder responses to LPS (van den Bos *et al*., unpublished data). This is not unprecedented as this has also been observed in mice: enhanced levels of IL1 are associated with a milder response to LPS (Alves-Rosa *et al*., 2002). It has been shown that following hatching *il1β* expression increases due to exposure to microbes in the medium (Galindo-Villegas *et al*., 2012). This tunes the activity of the innate immune system of the zebrafish larvae and determines thereby their disease resistance. In the laboratory however this may lead to variation between experimental series as the microbial load of the medium may vary from experimental series to series. It is clear that this warrants further studies. It has been suggested that tuning the innate immune system occurs through chromatin modifications (see e.g. Foster *et al*., 2007; Galindo-Vargas *et al*., 2012; Netea *et al*., 2016, 2017). Our future studies are directed at understanding the variable responses in this context.

The immune system affects HPI-axis activity and *vice versa* (Wendelaar Bonga, 1997). However, we did not address changes in HPI-axis activity as a consequence of our procedures (tail fin amputation or LPS exposure) in this study as the primary aim was to study whether our pre-treatments would affect immune function. Still changes in HPI-axis activity may be anticipated. For example, a recent study showed in European sea bass (*Dicentrarchus labrax*) larvae 5 day post-hatching that at 120 hours following infection with the bacterium *Vibrio anguillarum*, when mortality was already high, HPI-axis activity increased (Reyes-López *et al*., 2018). This increased activity coincided with increased expression of pro-inflammatory and anti-inflammatory genes. While the underlying mechanism was not clear as yet, the data show that infections may impact HPI-axis activity. It has been shown that early-life exposure to cortisol dampens the response of the HPI-axis to stressors in zebrafish larvae (Nesan and Vijayan, 2016). Hence, future studies should address how early-life exposure to cortisol affects the relationship between immune function and HPI-axis activity, and how this relates to the increased survival that we have observed here.

We have used 0-6 hpf cortisol exposure by the medium as a model or proxy of increased oocyte levels of cortisol due to chronic stress in mothers (van den Bos *et al*., 2019), while others have used micro-injection of cortisol in the yolk of single-cell embryos (Best *et al*., 2017; Nesan and Vijayan, 2012, 2016). Chronic stress, whether due to excessive predation, food shortage, crowding or out-of-range-temperatures, is associated with increased levels of cortisol (Wendelaar Bonga, 1997) and at this level these procedures may be a valid approach of mimicking maternal stress. Still, these different stressors may have additional neuro-endocrine and/or metabolic signatures that affect oocyte yolk-sac contents and hence thereby development of embryos and larvae. Thus, future experiments should compare the current results to data from chronically stressed mothers using different types of stressors.

### 4.5 Conclusion

These data show that early-life exposure to cortisol, as a model or proxy of maternal stress, induces an adaptive response to immune challenges, which seems mediated via the glucocorticoid receptor. These data are of relevance for both ecological research (Sopinka *et al*., 2017) and biomedical research (Stewart *et al*., 2014) in understanding the effects of stressful conditions and exposure to endocrine disruptors on disease susceptibility and survival of offspring.

## Abbreviations used in the manuscript

ANOVA: analysis of variance
GR: glucocorticoid receptor
dpf: days post-fertilisation
GFP: green fluorescent protein
hpa: hours post-amputation
hpf: hours post-fertilisation
HPI-axis: hypothalamus – pituitary - interrenal axis
IL: interleukine
KMO: Kaiser-Meyer-Olkin
LPS: lipopolysaccharide
MANOVA: multivariate analysis of variance
MR: mineralocorticoid receptor
ns: not significant
PBS: phosphate buffered saline
qPCR: quantitative polymerase chain reaction
SEM: standard error of the mean
TLR: Toll-like receptor
Tukey HSD: Tukey Honestly Significant Difference

## 5. Acknowledgements

We acknowledge Tom Spanings, Antoon van der Horst and Jeroen Boerrigter (Radboud University) and Guus van der Velden, Ulrike Nehrdich and Ruth van Koppen (Leiden University) for excellent fish care.

## 6. Conflict of interest

The authors have no conflict of interest to report.

## 7. Funding

The authors received no specific funding for this research.

## 8. Contributions

RvdB and MS conceived the project. SC, KT, JA and RW performed experiments. RvdB, SC, KT, JA and RW analysed data. JZ and SC performed qPCR analysis. RvdB, GF and MS wrote the manuscript.

## 10. Legends figures

**Supplementary figure 1.**
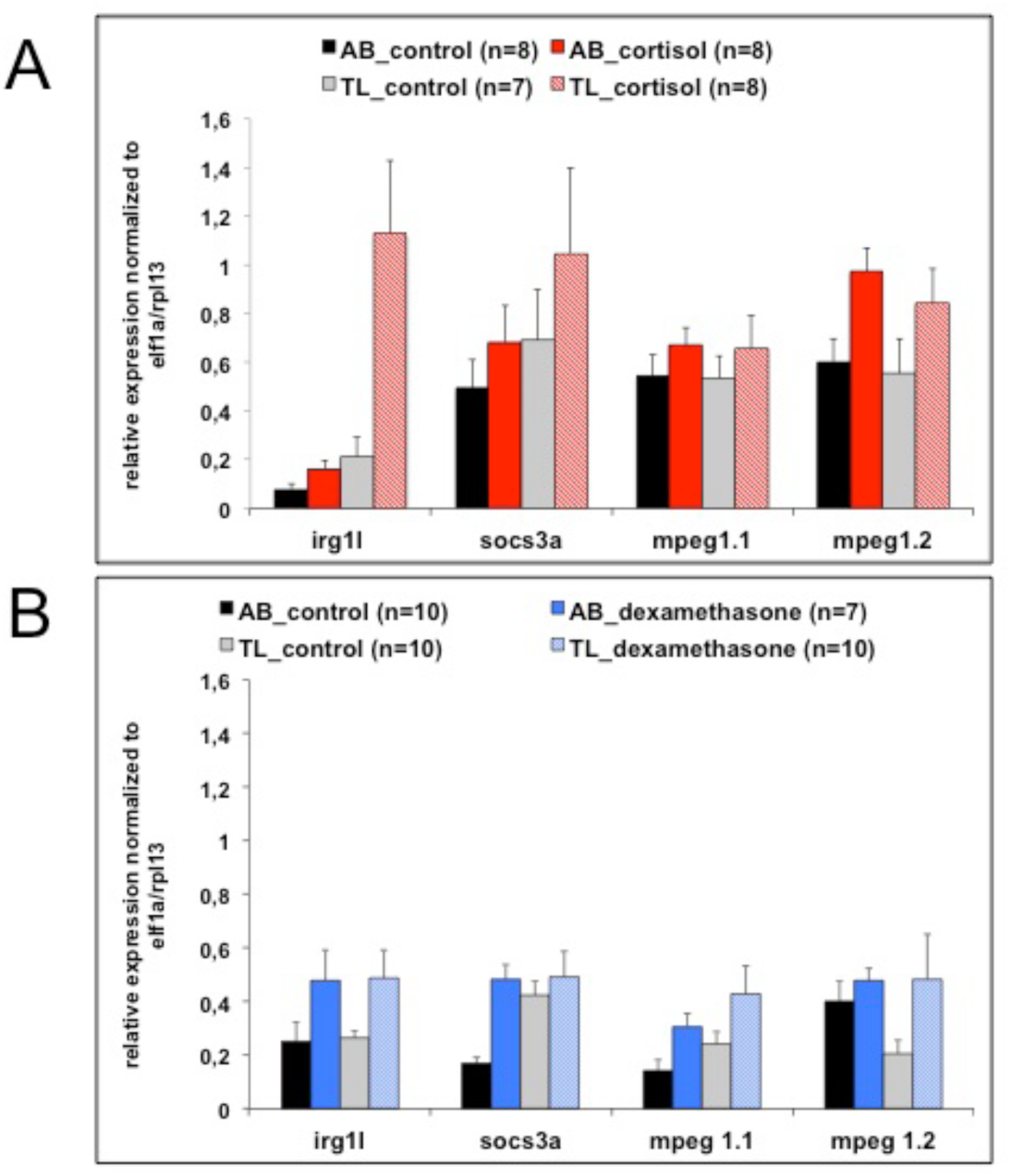
**panel A**: Transcript abundance (relative normalized expression; mean+SEM) of the different immune related genes for the different treatments (cortisol or control) and strains (AB or TL). One subject was removed from the statistical analyses (TL control) as it was a consistent outlier following Grubb’s outlier test. Treatment effects were found for *irg1l* (F(1,23)=11.789, p≤0.01; treatment * strain: F(1,23)=8.446, p≤0.01; AB: p≤0.05; TL: p≤0.01) and mpeg1.1 (F(1,23)=8.614, p≤0.01). **panel B**: Transcript abundance (relative normalized expression; mean+SEM) of the different immune related genes for the different treatments (dexamethasone or control) and strains (AB or TL). Treatment effects were found for *irg1l* (F(1,33)=6.484, p≤0.05), *socs3a* (F(1,33)=7.655, p≤0.01) and mpeg1.1 (F(1,33)=5.487, p≤0.05).

## Notes

#### Summary of Updates

One figure had been added (Figure 1: experiments); quality of figures has been improved; a few sections have been added to the limitations (line 614-636).

