## Supplementary material for "Early life glucocorticoid exposure modulates immune function in zebrafish (*Danio rerio*) larvae": van den Bos et al_Suppl_Table 1 (from another version of this preprint: version 1, 2019-12-12)

| Treatment | shape | | tail fin | | heart |
|---|---|---|---|---|---|
| | %curved | %normal | %swollen | %damaged | %oedema |
| **control (n=16)** | 50.0 | 18.8 | 75.0 | 6.3 | 6.3 |
| **cortisol (=n=19)** | 10.5 | 31.6 | 68.4 | 0.0 | 5.3 |
| **dexamethasone (n=18)** | 11.1 | 33.3 | 61.1 | 5.6 | 5.6 |
| **Chi-square** | 9.8, df=2, p≤0.0074 | | 2.12, df=4, ns | | 0.02, df=2, ns |

**Supplementary table 1**: Per cent of affected larvae at 4hrs following LPS exposure (0-0.5hr) for the different treatment groups: 0-6 hpf treatment with cortisol, dexamethasone or control medium. Shown are shape: curved; tail fin: swollen or damaged; heart: oedema. In the lower row overall statistics are indicated. It should be noted that both in the cortisol-treated (Chi-square=4.84, df=2, p≤0.03) and dexamethasone-treated (Chi-square=4.44, df=2, p≤0.04) groups fewer curved larvae were present than in the control-treated larvae.
