## Supplementary material for "Early life glucocorticoid exposure modulates immune function in zebrafish (*Danio rerio*) larvae": van den Bos et al_Suppl_Table 2 (from another version of this preprint: version 1, 2019-12-12)

| treatment | 0hr | 0.5hr | 1hr | 3hrs | 6hrs | 24hrs |
|---|---|---|---|---|---|---|
| **control** | 0 (24) | 0 (24) | 12.5 (24) | 0 (23) | 0 (21) | 15.8 (19) |
| **cortisol** | 0 (24) | 0 (24) | 0 (24) | 0 (24) | 0 (24) | 8.3 (24) |
| **dexamethasone** | 0 (24) | 0 (24) | 4.2 (24) | 4.3 (23) | 9.1 (22) | 9.1 (22) |

**Supplementary table 2**: Per cent of larvae, which are curved following LPS exposure (0-0.5hr), in the different treatment groups: 0-6 hpf treatment with cortisol-containing medium, dexamethasone-containing medium or control medium. Between brackets: the number of subjects at each time-point; these numbers vary over time as subjects die because of the LPS treatment.
