## Supplementary material for "Early life glucocorticoid exposure modulates immune function in zebrafish (*Danio rerio*) larvae": van den Bos et al_Suppl_Table 3 (from another version of this preprint: version 1, 2019-12-12)

| **Time (hrs)** | **component 1** | | | **component 2** | | | **component 3** | | |
|---|---|---|---|---|---|---|---|---|---|
| | **control** | **cortisol** | **dex** | **control** | **cortisol** | **dex** | **control** | **cortisol** | **dex** |
| **0** | +0.33±0.24 | +0.89±0.82 | +0.96±0.28 | -0.90±0.06 | -0.97±0.18 | -1.01±0.08 | +0.26±0.11$^{A,B}$ | +0.94±0.27$^{A}$ | -0.18±0.08$^{B}$ |
| **0.5** | -0.85±0.07 | -0.17±0.39 | -0.41±0.49 | -0.23±0.07$^{A}$ | -0.04±0.20$^{A}$ | -0.85±0.10$^{B}$ | -0.38±0.17$^{A}$ | +1.78±0.48$^{B}$ | -1.04±0.32$^{A}$ |
| **1** | -0.93±0.05 | -1.04±0.22 | -0.55±0.26 | +0.64±0.10$^{A}$ | -0.28±0.08$^{B}$ | +0.60±0.16$^{A}$ | +1.02±0.28 | -0.15±0.41 | -0.47±0.48 |
| **3** | +0.76±0.5$^{A}$ | -0.35±0.07$^{B}$ | +1.37±0.25$^{C}$ | +2.43±0.41$^{A}$ | +0.10±0.26$^{B}$ | +0.49±0.10$^{B}$ | -0.49±0.40 | -0.57±0.38 | -0.72±0.36 |

**Supplementary Table 3**: Mean (±SEM) factor regression scores for the different treatments of the 3 components that emerged from the PCA (see text for details). dex = dexamethasone. Super-scripts: different capitals indicate significant differences between groups (Tukey HSD) following a significant treatment effect at that particular time-point.
