## Supplementary material for "Early life glucocorticoid exposure modulates immune function in zebrafish (*Danio rerio*) larvae": van den Bos et al_Suppl_Table 4 (from another version of this preprint: version 1, 2019-12-12)

**Supplementary Table 4.** Mean (±SEM) transcript abundance of different genes involved in the inflammatory response (n=4 per time point). Numbers before genes indicate to which component of the PCA genes belong (see Supplementary Table 3 and text). Component 1: 'Toll-like receptor' genes; component 2: genes involved in LPS-induced inflammation; component 3: genes expressing mediators of the LPS-induced inflammatory response. dex = dexamethasone. Super-scripts: different capitals indicate significant differences between groups (Tukey HSD) following a significant treatment effect at that particular time-point.

MANOVA: treatment: Wilk's λ =0.014 $(F_{28,46})=12.431$, $p<0.001$); time: Wilk's λ =0.001 $(F_{42,68.994})=15.173$, $p<0.001$); treatment*time: Wilk's λ =0.001 $(F_{84,134.571})=4.549$, $p<0.001$)

| | **T = 0 hr** | | | **T = 0.5 hr** | | |
|---|---|---|---|---|---|---|
| **Gene** | **control** | **cortisol** | **dex** | **control** | **cortisol** | **dex** |
| 1 *tlr2* | 0.93±0.03$^{A}$ | 0.77±0.16$^{A}$ | 1.52±0.14$^{B}$ | 0.71±0.09 | 0.70±0.09 | 1.05±0.41 |
| 1 *tlr4ba* | 1.10±0.07 | 1.26±0.28 | 0.92±0.12 | 0.45±0.04$^{A,B}$ | 0.75±0.12$^{B}$ | 0.34±0.05$^{A}$ |
| 1 *tlr4bb* | 1.10±0.28 | 1.89±1.01 | 0.76±0.14 | 0.68±0.06 | 0.74±0.11 | 0.90±0.38 |
| 1 *tlr5a* | 0.76±0.06$^{A}$ | 0.88±0.09$^{A,B}$ | 1.19±0.14$^{B}$ | 0.59±0.05 | 0.72±0.10 | 0.56±0.06 |
| 1 *tlr5b* | 0.57±0.05 | 0.73±0.09 | 0.66±0.07 | 0.45±0.01$^{A}$ | 0.72±0.10$^{B}$ | 0.51±0.04$^{A,B}$ |
| 2 *myd88* | 0.53±0.06$^{A}$ | 0.38±0.02$^{B}$ | 0.23±0.01$^{C}$ | 0.64±0.03$^{A}$ | 0.52±0.04$^{A}$ | 0.23±0.03$^{B}$ |
| 2 *il1β* | 0.016±0.002 | 0.018±0.004 | 0.020±0.001 | 0.15±0.01 | 0.22±0.02 | 0.10±0.02 |
| 2 *il10* | 0.47±0.07 | 0.43±0.11 | 0.42±0.03 | 0.36±0.03 | 0.51±0.18 | 0.28±0.06 |
| 3 *cxcr4a* | 0.51±0.04 | 0.66±0.05 | 0.60±0.07 | 0.51±0.03$^{A}$ | 0.87±0.14$^{B}$ | 0.36±0.02$^{A}$ |
| 3 *cxcr4b* | 0.38±0.03$^{A}$ | 0.56±0.04$^{B}$ | 0.44±0.04$^{A,B}$ | 0.36±0.05$^{A}$ | 0.75±0.09$^{B}$ | 0.41±0.03$^{A}$ |
| 3 *ptpn6* | 0.83±0.05$^{A}$ | 0.77±0.07$^{A,B}$ | 0.62±0.02$^{B}$ | 0.58±0.02$^{A,B}$ | 0.80±0.08$^{A}$ | 0.47±0.07$^{B}$ |

| | **T = 1 hr** | | | **T = 3 hrs** | | |
|---|---|---|---|---|---|---|
| **Gene** | **control** | **cortisol** | **dex** | **control** | **cortisol** | **dex** |
| 1 *tlr2* | 0.62±0.07 | 0.92±0.14 | 0.96±0.25 | 1.14±0.06$^{A,B}$ | 0.98±0.11$^{A}$ | 1.54±0.14$^{B}$ |
| 1 *tlr4ba* | 0.38±0.03 | 0.31±0.06 | 0.43±0.11 | 0.29±0.03$^{A}$ | 0.34±0.04$^{A}$ | 0.72±0.06$^{B}$ |
| 1 *tlr4bb* | 0.79±0.21$^{A}$ | 0.23±0.06$^{B}$ | 0.73±0.12$^{A.B}$ | 1.89±0.23$^{A}$ | 0.60±0.10$^{B}$ | 1.34±0.12$^{A}$ |
| 1 *tlr5a* | 0.66±0.04 | 0.63±0.07 | 0.74±0.12 | 1.19±0.09 | 0.94±0.12 | 1.19±0.14 |
| 1 *tlr5b* | 0.39±0.02$^{A,B}$ | 0.33±0.04$^{B}$ | 0.46±0.04$^{A}$ | 0.80±0.08$^{A}$ | 0.50±0.04$^{B}$ | 0.99±0.02$^{A}$ |
| 2 *myd88* | 0.66±0.10$^{A,B}$ | 0.41±0.03$^{A}$ | 1.00±0.12$^{B}$ | 1.08±0.14$^{A}$ | 0.48±0.08$^{B}$ | 0.89±0.05$^{A}$ |
| 2 *il1β* | 0.54±0.05 | 0.45±0.03 | 0.52±0.07 | 1.25±0.22$^{A}$ | 0.48±0.08$^{B}$ | 0.64±0.06$^{B}$ |
| 2 *il10* | 0.69±0.03$^{A}$ | 0.36±0.05$^{B}$ | 0.46±0.06$^{B}$ | 1.51±0.18$^{A}$ | 0.46±0.05$^{B}$ | 0.65±0.07$^{B}$ |
| 3 *cxcr4a* | 0.73±0.09$^{A}$ | 0.40±0.05$^{B}$ | 0.48±0.09$^{A,B}$ | 0.68±0.10$^{A}$ | 0.48±0.07$^{A,B}$ | 0.39±0.04$^{B}$ |
| 3 *cxcr4b* | 0.66±0.08 | 0.44±0.06 | 0.44±0.06 | 0.29±0.09 | 0.46±0.05 | 0.47±0.07 |
| 3 *ptpn6* | 0.71±0.03 | 0.76±0.12 | 0.47±0.08 | 0.42±0.04 | 0.47±0.05 | 0.44±0.07 |

| | **Statistics: two-way ANOVA** | | | | |
|---|---|---|---|---|---|
| **Gene** | **Treatment F(2,36)** | **Time F(3,36)** | **Treatment * Time F(6,36)** | **Tukey HSD Treatment** | **Tukey HSD Time** |
| 1 *tlr2* | **8.171;p≤0.001** | **3.938,p≤0.016** | 0.876;ns | con$^{A}$ cort$^{A}$ dex$^{B}$ | 0$^{A,B}$ 30$^{A}$ 60$^{A}$ 180$^{B}$ |
| 1 *tlr4ba* | 1.026;ns | **29.421;p<0.001** | **3.654;p≤0.006** | | 0$^{A}$ 30$^{B}$ 60$^{B}$ 180$^{B}$ |
| 1 *tlr4bb* | 0.575;ns | **3.128;p≤0.04** | *2.262;p≤0.06* | | |
| 1 *tlr5a* | *2.883;p≤0.07* | **19.207;p<0.001** | **2.382;p≤0.048** | | 0$^{A}$ 30$^{B}$ 60$^{B}$ 180$^{A}$ |

| | | | | | |
|---|---|---|---|---|---|
| 1 *tlr5b* | **3.846;p≤0.03** | **23.574;p<0.001** | **8.596;p<0.001** | con$^{A}$ cort$^{A,B}$ dex$^{B}$ | 0$^{A,B}$ 30$^{A}$ 60$^{C}$ 180$^{B}$ |
| 2 *myd88* | **16.301;p<0.001** | **25.213;p<0.001** | **11.970;p<0.001** | con$^{A}$ cort$^{B}$ dex$^{C}$ | 0$^{A}$ 30$^{A}$ 60$^{B}$ 180$^{B}$ |
| 2 *il1β* | **7.302;p≤0.002** | **59.251;p<0.001** | **6.794;p<0.001** | con$^{A}$ cort$^{B}$ dex$^{B}$ | 0$^{A}$ 30$^{A}$ 60$^{B}$ 180$^{C}$ |
| 2 *il10* | **16.349;p<0.001** | **18.287;p<0.001** | **9.450;p<0.001** | con$^{A}$ cort$^{B}$ dex$^{B}$ | 0$^{A}$ 30$^{A}$ 60$^{A}$ 180$^{B}$ |
| 3 *cxcr4a* | **5.776;p≤0.007** | 0.757;ns | **6.319;p<0.001** | con$^{A}$ cort$^{A}$ dex$^{B}$ | |
| 3 *cxcr4b* | **6.701;p≤0.003** | *2,631;p≤0.07* | **5.260;p≤0.001** | con$^{A}$ cort$^{B}$ dex$^{A}$ | |
| 3 *ptpn6* | **10.666;p<0.001** | **11.648,p<0.001** | *1.958;p≤0.098* | con$^{A}$ cort$^{A}$ dex$^{B}$ | 0$^{A}$ 30$^{A}$ 60$^{A}$ 180$^{B}$ |
